# Primed to Die: An Investigation of the Genetic Mechanisms Underlying Noise-Induced Hearing Loss and Cochlear Damage in Homozygous Foxo3-knockout Mice

**DOI:** 10.1101/2021.03.12.435183

**Authors:** Holly J. Beaulac, Felicia Gilels, Jingyuan Zhang, Sarah Jeoung, Patricia M. White

**Affiliations:** Department of Neuroscience, Ernest J. Del Monte Institute for Neuroscience, University of Rochester School of Medicine and Dentistry, Rochester, NY, USA; University of Rochester School of Medicine and Dentistry, Rochester, NY, USA

**Author notes:** Now at Department of Pathology, University of Rochester School of Medicine and Dentistry, Rochester, NY, USA. Now at Department of Otolaryngology, Otolaryngology-Head and Neck Surgery, Harvard Medical School, Boston Children’s Hospital Center for Life Science, Boston, MA, USA. Corresponding Author. Department of Neuroscience, Ernest J. Del Monte Institute for Neuroscience, University of Rochester School of Medicine and Dentistry, 601 Elmwood Avenue, Rochester, NY, 14642, USA. **Abbreviations.** NIHL, noise-induced hearing loss; OHC, outer hair cell; IHC, inner hair cell; SC, supporting cell; P, postnatal day; WT, wild-type; ABR, auditory brainstem response; DPOAE, distortion product otoacoustic emission; DPN, days post noise; HPN, hours post noise; kHz, kilohertz; dB, decibel; DEGs, differentially expressed genes; GDPD3, Glycerophosphodiester Phosphodiesterase Domain Containing 3; LPA, lysophosphatidic acid.

## Abstract

The prevalence of noise-induced hearing loss (NIHL) continues to increase, with limited therapies available for individuals with cochlear damage. We have previously established that the transcription factor FOXO3 is necessary to preserve outer hair cells (OHCs) and hearing thresholds up to two weeks following a mild noise exposure in mice. The mechanisms by which FOXO3 preserves cochlear cells and function are unknown. In this study, we analyzed the immediate effects of mild noise exposure on wild-type, *Foxo3* heterozygous (*Foxo3^+/KO^*), and *Foxo3* knock-out (*Foxo3^KO/KO^*) mice to better understand FOXO3’s role(s) in the mammalian cochlea. We used confocal and multiphoton microscopy to examine well-characterized components of noise-induced damage including calcium regulators, oxidative stress, necrosis, and caspase-dependent and -independent apoptosis. Lower immunoreactivity of the calcium buffer oncomodulin in *Foxo3^KO/KO^* OHCs correlated with cell loss beginning 4 hours post-noise exposure. Using immunohistochemistry, we identified parthanatos as the cell death pathway for OHCs. Oxidative stress response pathways were not significantly altered in FOXO3’s absence. We used RNA sequencing to identify and RT-qPCR to confirm differentially expressed genes. We further investigated a gene downregulated in the unexposed *Foxo3^KO/KO^* mice that may contribute to OHC noise susceptibility. Glycerophosphodiester Phosphodiesterase Domain Containing 3 (GDPD3), a possible endogenous source of lysophosphatidic acid (LPA), has not previously been described in the cochlea. As LPA reduces OHC loss after severe noise exposure, we treated noise exposed *Foxo3^KO/KO^* mice with exogenous LPA. LPA treatment delayed immediate damage to OHCs but was insufficient to ultimately prevent their death or prevent hearing loss. These results suggest that FOXO3 acts prior to acoustic insult to maintain cochlear resilience, possibly through sustaining endogenous LPA levels.

## Introduction

Noise induced hearing loss (NIHL) is a pervasive health threat. In 2017, the NIDCD estimated that despite increased awareness and access to protective equipment, nearly 40 million American adults had signs of NIHL^1^. Therapies for NIHL remain limited, in many cases due to the permanent loss of cochlear sensory cells. Individuals have varying susceptibilities to noise damage, partially due to differences in gene expression crucial for hearing recovery and cochlear preservation^2^. By studying the underlying genetic mechanisms of NIHL susceptibility, our goal is to better prevent or mitigate NIHL using biological approaches.

Transcriptional regulators comprise a class of NIHL susceptibility genes, including forkhead box O-3 (FOXO3)^3^. FOXO3 is a winged helix/forkhead class transcription factor involved in cellular processes including autophagy^4^, survival^5^, stress resistance^6–9^, apoptosis^10–12^, and longevity^13–16^. Individuals may harbor genetic variants that can lead to varying levels of FOXO3 transcription, with its spatiotemporal regulatory roles susceptible to manipulation in autoimmune diseases^17–19^ and cancers^20,21^. FOXO3 is expressed from birth into adulthood throughout the mammalian cochlea including inner and outer hair cells (IHCs and OHCs), supporting cells (SCs), and spiral ganglion neurons (SGNs)^22^. FOXO3 is capable of translocating from the cytoplasm to nucleus in response to a stressor, as SGNs respond to non-traumatic noise^22^. This suggests a role in driving gene transcription, which could prevent additional damage or promote death in dysfunctional cells. We have previously shown that FOXO3 is required to preserve OHCs and hearing thresholds following mild noise exposure^23^. In the hematopoietic system, loss of FOXO3 leads to the overaccumulation of reactive oxygen species (ROS), leading to the death of hematopoietic stem cells^24^ and developing erythrocytes^25^. Accumulation of ROS in cochlear cells after noise exposure also drives cell death^26–28^. We hypothesize that in the absence of FOXO3 following noise exposure, higher levels of reactive oxygen species would accumulate in *Foxo3^KO/KO^* OHCs versus WT OHCs, triggering an apoptotic cascade.

To better understand FOXO3’s role in NIHL susceptibility, we analyzed the immediate effects of mild noise exposure on wild-type (WT, *Foxo3^+/+^*), *Foxo3* heterozygous (*Foxo3^+/KO^*), and *Foxo3* knock-out (*Foxo3^KO/KO^*) mice. Previous studies have shown that after traumatic noise exposure, OHC damage drives apoptosis, necroptosis, and a third death pathway^29–31^. Depending on the severity of the traumatic exposure, OHC caspase-3 activation may be immediately evident^29^, or it may develop over the following 24 hours post noise (HPN)^32^. OHC loss continues over a period of days to weeks^32,33^. The degree of OHC loss seen in the *Foxo3^KO/KO^* cochlea after a mild noise exposure could reflect either changes in cochlear homeostasis prior to damage or a compromised damage response. Determining the time course of OHC death could shed light on when FOXO3 acts to protect the cochlea.

To this end, we characterized the time course of OHC loss from noise in the *Foxo3^KO/KO^* mouse and identified a likely mode of cell death. We compared activation of known damage pathways in the *Foxo3^KO/KO^* cochlea to that of the WT cochlea to assess their involvement. We performed RNA-SEQ, identified a candidate downstream effector enzyme, and tested its involvement by exogenous supplementation of its product. Our goals were to determine what pathways were immediately active after noise and when therapeutic intervention would be most efficacious to mitigate the effects of FOXO3’s absence.

## Results

### Establishing the time course of OHC loss in the *Foxo3^KO/KO^* noise-exposed cochlea

*Foxo3^KO/KO^* mice acquire severe permanent threshold shifts throughout the cochlea after mild noise exposure^23^. By two weeks, about one third of *Foxo3^KO/KO^* OHCs were lost from the basal cochlea without any IHC loss^23^. WT littermates experienced only temporary threshold shifts and no cell loss from the same exposure^23^. These observations did not establish when or how *Foxo3^KO/KO^* OHCs are lost. Here we characterize cellular pathology in *Foxo3^KO/KO^* and WT cochleae immediately after noise exposure. We used cochlear cryosectioning to obtain cross-sections for detailed cellular analysis (Fig. S1A), and whole mounts preparations for cellular quantification according to tonotopic frequency (Fig. S1B). The latter is important as high frequency (basal) OHCs are more susceptible to trauma^34–36^. To test for haploinsufficiency, we analyzed the FOXO3 heterozygote (*Foxo3^+/KO^)*. We measured *Foxo3^+/KO^* auditory brainstem responses (ABRs) and distortion product otoacoustic emissions (DPOAEs) at baseline, 1-day post-noise (1 DPN), and 14 DPN (Fig. S2). Since *Foxo3^+/KO^* mice displayed temporary threshold shifts and recovery similar to WT mice^23^, one *Foxo3* allele appears sufficient for cochlear homeostasis.

To establish the time course of OHC loss in *Foxo3^KO/KO^* mice, we exposed adult WT, *Foxo3^+/KO^*, and *Foxo3^KO/KO^* mice to mild noise (Fig. 1A). Cochleae were isolated at 0.5, 4, and 24 HPN, and fluorescent antibodies for oncomodulin (OCM) and myosin 7a (MYO7a)^37^ were used to quantify OHCs. OCM is an EF-hand calcium binding protein highly expressed in OHCs^38^, and its deletion confers progressive hearing loss^39^. Representative 24 kHz (middle) regions of *Foxo3^KO/KO^* cochleae are presented in Fig. 1B-E’. Under baseline conditions, OCM and MYO7a were co-expressed within the OHCs (Fig. 1B-B’). OCM expression was reduced in some *Foxo3^KO/KO^* MYO7a+ OHCs after 4 HPN (Fig. 1D-D’, yellow arrows), becoming more apparent at 24 HPN in rows 2 and 3 (Fig. 1E-E’, yellow arrows).

**Figure 1.**
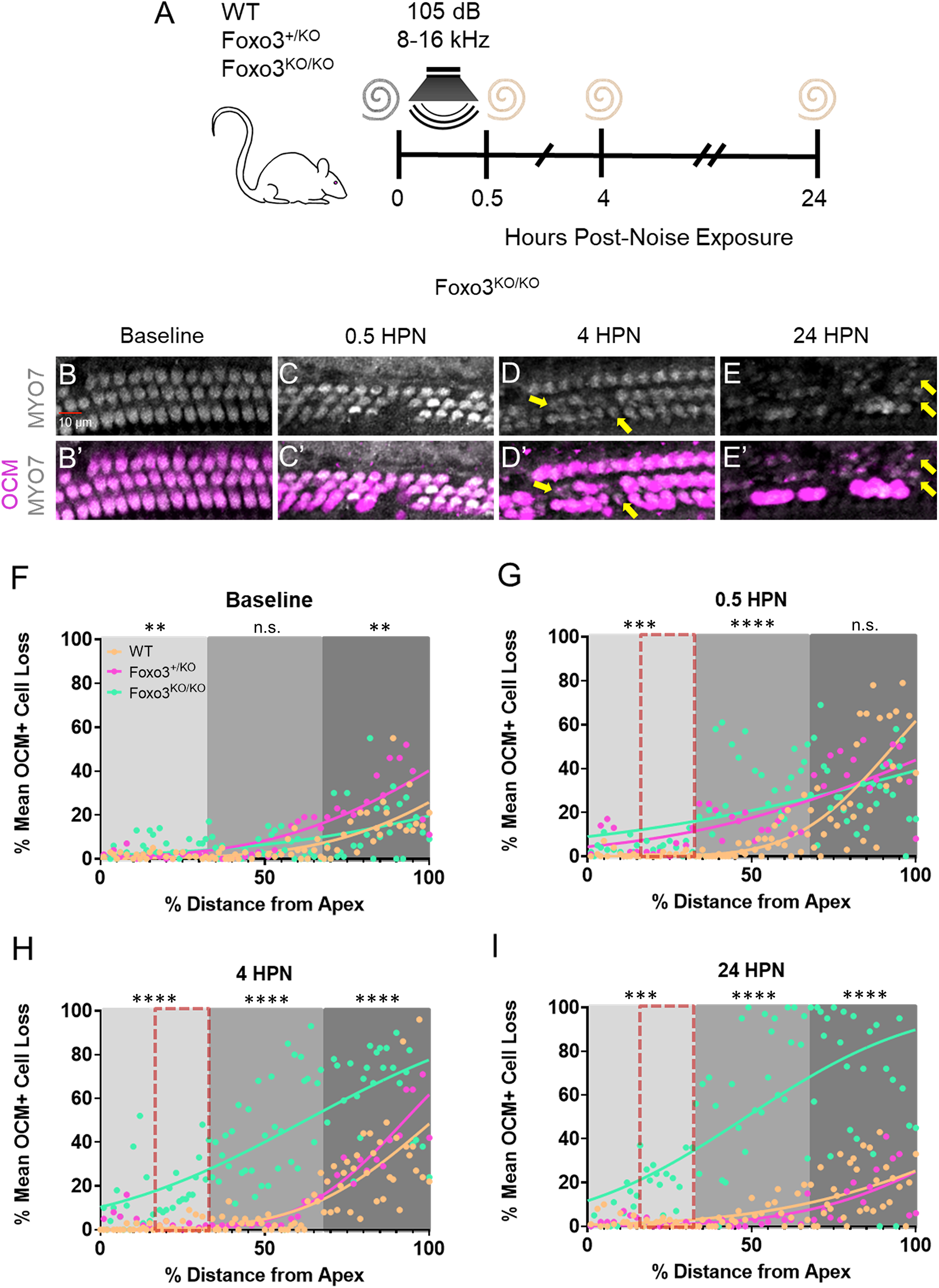
OCM modulation observed immediately following noise exposure. **A)** Noise exposure method for WT, *Foxo3^+/KO^*, and *Foxo3^KO/KO^* mice. Cochleae were extracted for whole mount immunohistochemistry immediately following the exposure (0.5 hours post-noise exposure or HPN), at 4 HPN, or 24 HPN. Unexposed littermates served as baseline controls. **B-E)** Expression of MYO7 (white) was maintained in the *Foxo3^KO/KO^* OHCs following noise exposure (yellow arrows). **B’-E’)** Over the same time course, OCM (pink) immunoreactivity decreased in OHCs, mainly in rows 2 and 3. n = 3, scale bar = 10 µm, 20x magnification, ∼24 kHz region. **F-I)** Cochleograms of % mean OCM+ cell loss (y-axis) plotted against % distance from the cochlear apex (x-axis). Each dot represents the mean number of OCM+ cells counted in 100 µM intervals along the length of cochleae (4-86 kHz) pooled (n = 3-6 per genotype/condition). Cochlear tonotopic regions are presented by background color: Apex = light gray, Middle = gray, Base = dark gray. Non-parametric interpolation lines are presented for each group: WT (orange), *Foxo3^+/KO^* (pink), and *Foxo3^KO/KO^* (green). **G-I)** The noise band is presented as a red dotted outline. Kruskal-Wallis rank sum test adjusted for multiple comparisons, alpha = 0.05, *p < 0.05, **p < 0.01, ***p < 0.001, ****p < 0.0001.

To determine the overall number of OCM+ OHCs after noise exposure, cochleae were stained with DAPI and antibodies against OCM, MYO7a or Cytochrome-C (CytC), a mitochondria-associated hemeprotein strongly expressed in OHCs^40^. Cochleograms were generated for each genotype and timepoint. All three genotypes displayed slightly more baseline OCM-low OHCs in the basal cochlear third (Fig. 1F). The *Foxo3^KO/KO^* apex and *Foxo3^+/KO^* base showed increased OCM loss (Fig. 1F; Apex p = 0.0042, Middle p = 0.3000, Base p = 0.0043, Kruskal-Wallis). Immediately following noise, loss of OCM+ OHCs increased in both *Foxo3^KO/KO^* and *Foxo3^+/KO^* cochleae at the apical (p = 0.0002) and middle (p < 0.0001) regions. All three genotypes had similar losses in the base (p = 0.3000, Fig. 1G; Kruskal-Wallis). By 4 HPN, loss of OCM+ cells increased along the entire lengths of *Foxo3^KO/KO^* cochleae (all turns p < 0.0001) while *Foxo3^+/KO^* and WT numbers recovered to baseline (Fig. 1H). *Foxo3^KO/KO^* cochleae had greater OCM+ cell loss by 24 HPN (all turns p < 0.0001) as the other two genotypes regained OCM immunoreactivity (Fig. 1I, Kruskal-Wallis). These data suggest that OCM dynamically modulates, and its normal response to noise exposure is exacerbated in the *Foxo3^KO/KO^* cochleae.

To assess cellular damage, we rendered the 12 (apical) and 24 (middle) kHz regions in Imaris (Fig. 1). We focus on 24 kHz WT and *Foxo3^KO/KO^* renderings (Fig. 2A-H”) and mean OHC counts of all three genotypes (Fig. 2I-K and Fig. S3). At baseline, CytC distribution in OHCs was similar to OCM (Fig. 2A”, E”). WT mice showed consistent OCM immunoreactivity at baseline and following noise exposure (Fig. 2A-D”). At baseline, some low OCM-expressing OHCs were detected in *Foxo3^+/KO^* mice (Fig. S3N), and *Foxo3^KO/KO^* OHCs were comparable to WT cells (Fig. 2E-E”). Immediately following noise exposure, many *Foxo3^KO/KO^* OHCs lost OCM immunoreactivity (Fig. 2F-F”, yellow arrows). By 4 HPN, pyknotic nuclei were present in OHC regions lacking OCM (Fig. 2G-G”). By 24 HPN, many OCM+/CytC+ cells were lost and fewer OHC nuclei remained in the *Foxo3^KO/KO^* (Fig. 2H-H”, white arrows). Most WT OHCs expressed OCM by 24 HPN (Fig. 2D-D”, yellow arrows). Notably, their nuclei were neither fragmented nor expanded. Although WT and *Foxo3^+/KO^* mice displayed fluctuations in OCM immunoreactivity, unlike the *Foxo3^KO/KO^* mice this did not resolve into OHC loss by 24 HPN (Fig. 2I, J p < 0.0001, two-way ANOVA). The 12 kHz region exhibited similar but less drastic OCM changes than the 24 kHz region, and *Foxo3^KO/KO^* OHCs were better preserved (Fig. S4). In the absence of FOXO3, weak OCM immunoreactivity lasting beyond 4 HPN correlated with OHC apoptosis.

**Figure 2.**
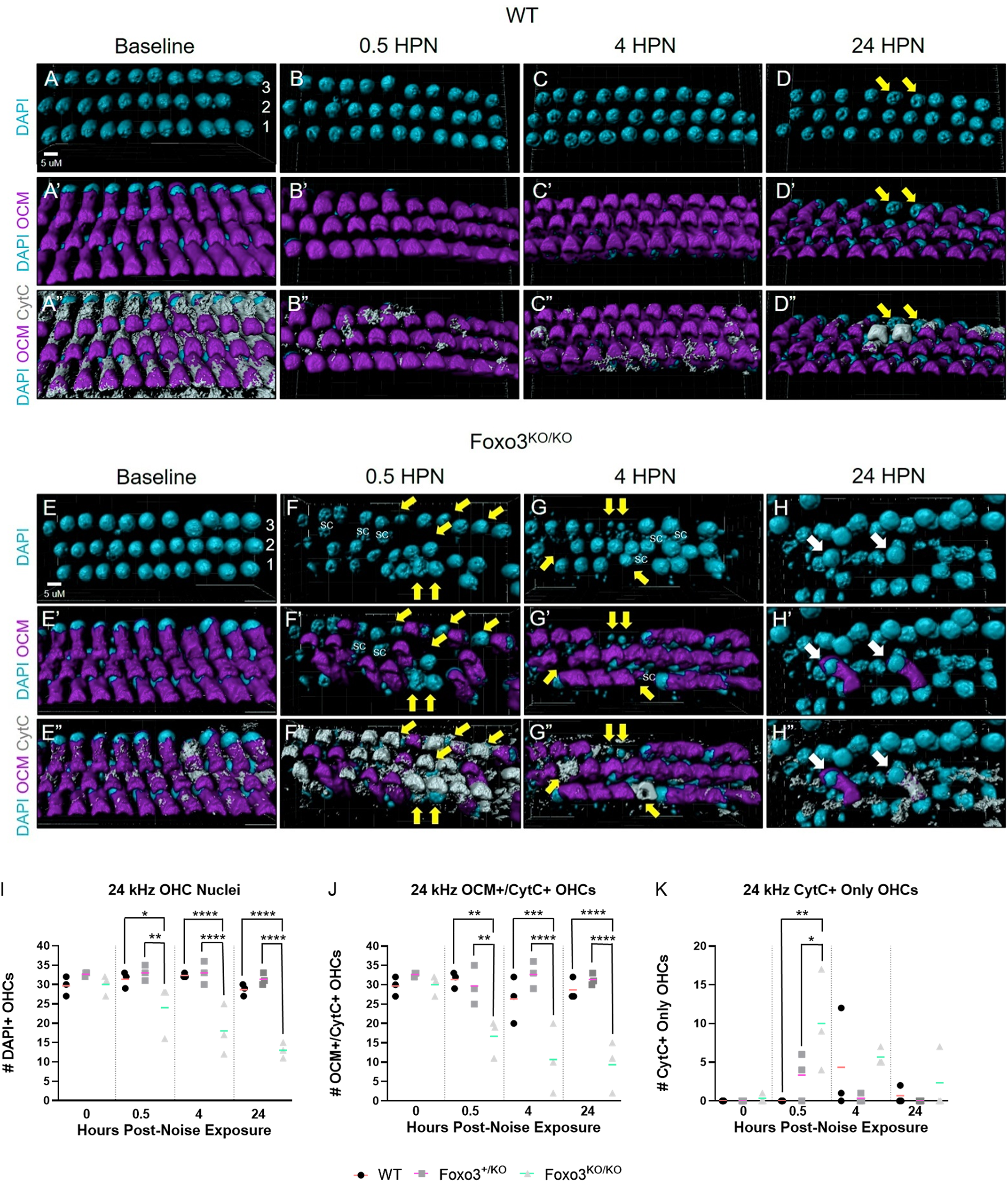
Loss of OCM immunoreactivity precedes apoptosis in *Foxo3^KO/KO^* OHCs. After immunostaining in whole mount, the 24 kHz cochlear region was mapped, imaged using confocal microscopy at 150x magnification, and rendered in Imaris (n = 3 cochleae per genotype/condition). **A-H)** The rendered images of OHCs colored cyan for DAPI+ cell nuclei. **A’-H’)** Renderings of DAPI+/OCM+ OHCs with OCM in magenta. **A”-H”)** Renderings of DAPI+/OCM+/CytC+ cells with CytC in gray. Scale bar = 5 µm. **A-C”)** Little to no change in OCM and CytC localization through 4 HPN in WT OHCs. **D-D”)** By 24 HPN, WT OHCs only expressing CytC+ (yellow arrows) are in the minority of cells likely damaged by the noise exposure. **F’, F”)** At 0.5 HPN, a large number of *Foxo3^KO/KO^* OHCs lose OCM immunoreactivity (yellow arrows). **G, G”)** By 4 HPN, several OHC regions with low OCM contain pyknotic or missing nuclei (yellow arrows). **H’, H”)** Only two OCM+/CytC+ OHCs remain in any of the standard three rows (white arrows); the other nuclei belong to SCs as they exist in a lower z-plane and have a larger mean diameter. **I-K)** OHC counts for all three genotypes expressing specific fluorophores (y-axis) are graphed versus the noise exposure timeline (x-axis). **I)** Numbers of DAPI+ OHC nuclei (y-axis) present across the time course (x-axis); supporting cell nuclei unable to be excluded in the renderings due to thresholding limitations were omitted. **J)** Counts of OHCs as calculated by totaling OCM+/CytC+ cells (y-axis). **K)** Counts of OHCs only expressing CytC (y-axis). WT (black circles, orange mean bar), *Foxo3^+/KO^* (gray squares, pink mean bar), and *Foxo3^KO/KO^* (light gray triangles, green mean bar) total cell counts per 1024 x 400 x 20-pixel selection. n = 3 per genotype/condition, two-way ANOVA with Tukey’s test for multiple comparisons, alpha = 0.05, *p < 0.05, **p < 0.01, ***p < 0.001, ****p < 0.0001.

### Caspase-independent parthanatos may drive OHC loss

FOXO3 regulates caspase-dependent apoptosis in many tissues by promoting transcription of Bax and Bim^10,11^. Nuclear fragmentation, indicating apoptosis, correlated with decreased OCM in *Foxo3^KO/KO^* OHCs following noise exposure. Activated CASP3 immunostaining was not observed (Fig. 3A-D’), suggesting a caspase-independent apoptotic pathway in noise-induced OHC death^31,41–44^. Parthanatos is a rapid caspase-independent apoptotic mechanism seen in cells primed for death^40^. It is identified by the nuclear localization of the mitochondrial protein AIFM1, which initiates a cell death program^45^. No nuclear AIFM1 was seen in WT OHCs (Fig. 3). We saw low AIFM1 immunoreactivity in the *Foxo3^KO/KO^* at baseline, which became elevated in OCM-low OHC nuclei at 0.5 HPN (Fig. 3H-H’”, yellow arrows). These results are consistent with parthanatos as a pathway for OHC death in *Foxo3^KO/KO^* cochleae.

**Figure 3.**
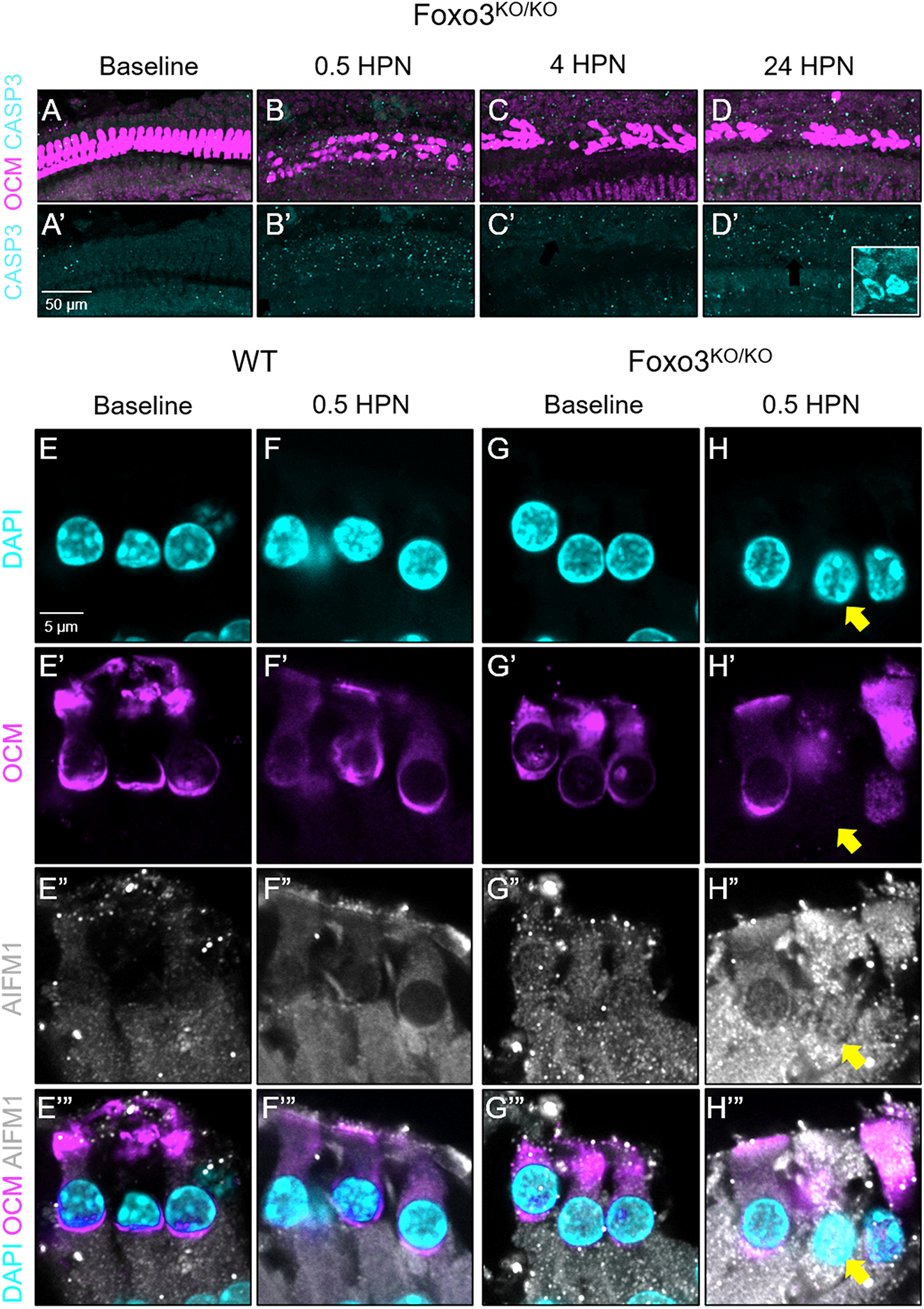
*Foxo3^KO/KO^* OHCs do not initiate caspase-dependent apoptosis but may activate parthanotic pathways after mild noise exposure. **A-D’)** Multiphoton images of the middle, approximately 24 kHz region, in whole-mounted cochleae over the established time course. Tissues were analyzed for OCM (magenta) and CASP3 (cyan). **D’)** Insert = positive control tissue: CASP3+ apoptotic SGNs after chronic cigarette smoke exposure. n = 3-5 per time point, 20x magnification, scale bar = 50 μm. **E-H’”)** Single-matched optical sections from confocal images of 24 kHz OHCs under baseline conditions and 0.5 HPN. WT **(E-F’”)** and *Foxo3^KO/KO^* (G-H’”) cochleae were sectioned and immunostained for detection of DAPI (cyan, **E-H**), OCM (magenta, **E’-H’**), and AIFM1 (white, **E”-H”**). **H-H’”)** The mitochondrial protein AIFM1, localized to the nucleus in the *Foxo3^KO/KO^* OCM-low OHC (yellow arrow), an indication of parthanatos. n = 4 per genotype/condition, 200x magnification, scale bar = 5 μm.

### Differential expression analysis of genes following noise exposure shows enhancement in calcium signaling and oxidative stress response pathways

Absence of the transcription factor FOXO3 could alter gene expression, leaving OHCs primed for death. *Foxo3^KO/KO^* and WT cochleae were processed at 0, 4, and 24 HPN for mRNA extraction (Fig. 4A). Differentially expressed genes (DEGs) between genotypes and time points were identified with RNA-sequencing. Gene ontology was used to perform enrichment analysis on DEGs. Calcium signaling, the oxidative stress response, and some cytokine signaling pathways were enhanced by 4 HPN in the *Foxo3^KO/KO^* (Fig. 4B). Changes in calcium signaling genes correlated with the OCM modulation observed shortly after noise. Clustering analysis for the *Foxo3^KO/KO^* versus WT 4 HPN comparison found 8 distinct gene clusters (Fig. 4C, Fig. S5). Several genes matched with the canonical pathways assigned at 4 HPN and are listed in Supplementary Table 3. Volcano plots enable visualization of DEG distribution (Fig. 4D-E). Notably, the genes Gdpd3, Ypel3, and Gatsl2 were downregulated at baseline and following noise exposure (Fig. 4D-F). We include these genes in our analysis of OHC-associated factors (Table 1). The transcription factor *JunB* was up-regulated (Log FC = 1.73, Log CPM = 5.21, p = 8.52E-16) in the *Foxo3^KO/KO^* cochleae at 4 HPN. This was confirmed with RT-qPCR (fold gene expression = 4.35). Jun family members JunB, cJUN, and JNK are associated with the oxidative stress response^46,47^. Provided these RNA sequencing results and known associations between FOXO3 and oxidative stress regulation in other tissues, we examined the oxidative stress response following noise.

**Figure 4.**
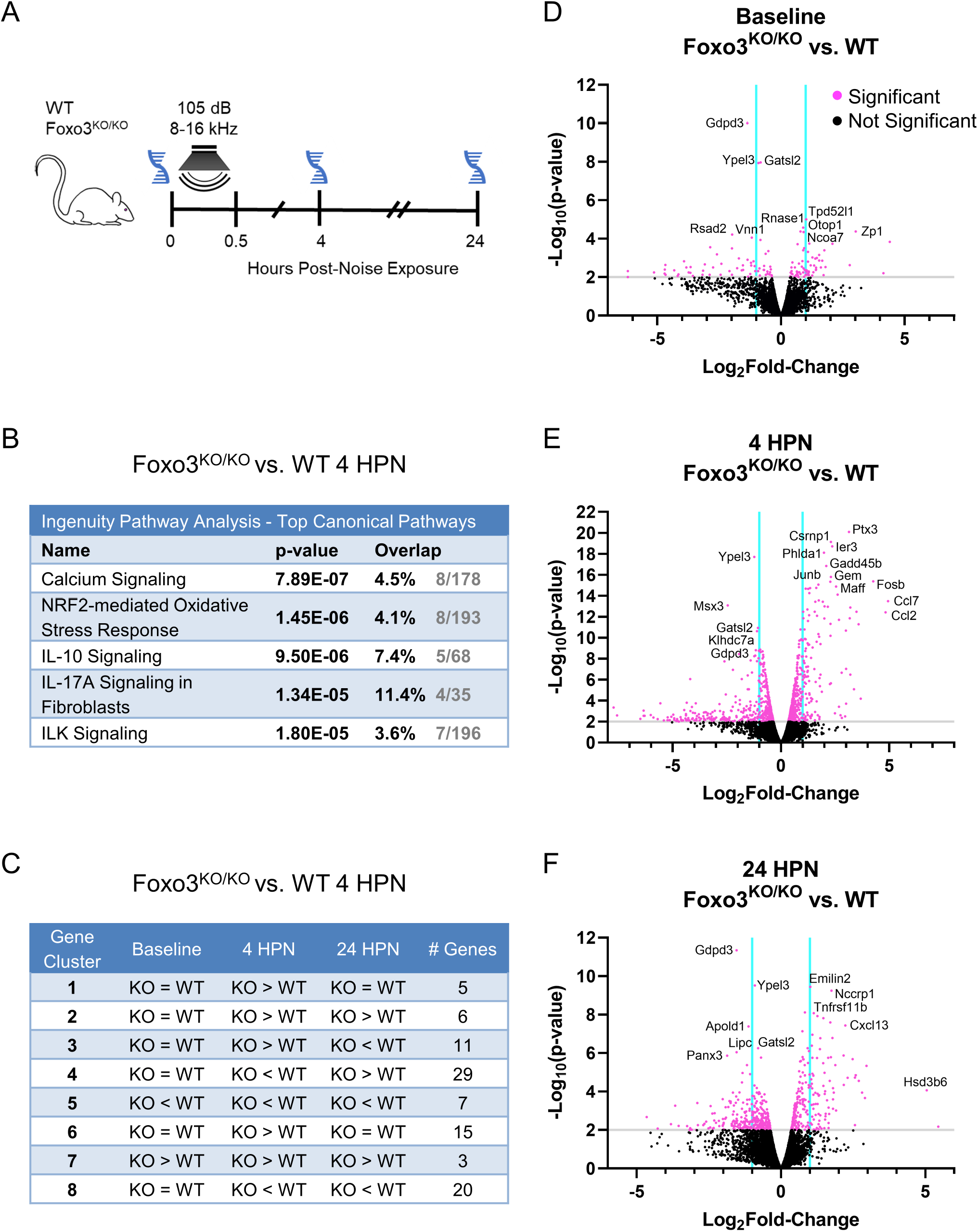
RNA sequencing revealed calcium signaling and the oxidative stress response as active pathways following noise exposure. **A)** *Foxo3^KO/KO^* and WT mice were subjected to noise and had their cochleae removed for mRNA extraction at 0, 4, and 24 HPN (n = 6-7 pooled cochleae per genotype/condition). **B)** Top canonical pathways in *Foxo3^KO/KO^* versus WT cochleae at 4 HPN using Ingenuity Gene Ontology. The p-value indicates the statistical significance of the overlapping genes within the canonical pathway. **C)** Gene expression clustering was performed for the *Foxo3^KO/KO^* versus WT at 4 HPN to produce 8 clusters with similarly behaving genes. Clustering traces and the detailed gene table are available in Supplements (Fig. S7, Table S3). **D-F)** Volcano plots of differentially expressed genes at baseline **(D)**, 4 HPN **(E)**, and 24 HPN **(F)**. The y-axis represents −log_10_(p-value) and x-axis is the log_2_fold-change. Significantly differentiated genes (-log_10_(p-value) ≥ 2) are denoted by pink dots above the gray line. Non-significant genes are represented by black dots. Cyan lines divide genes that fall between a −1 to 1 log fold-change.

**Table 1.** Curated changes in genes expressed in OHCs at baseline. Foxo3^KO/KO^ cochleae were compared to wild-type littermates in bulk RNA sequencing preparations. Genes expressed in OHCs were identified through comparison with the gEAR portal (He dataset). Differential expression was observed for some stereociliary and actin-binding protein mRNA (blue), as well as mRNA encoding hair cell-specific transcription/translation factors (pink). Transcripts for three stress response proteins were identified (yellow), along with those associated with neurotransmitter channels and receptors (green). Note that MYH7 encodes the myosin heavy chain beta isoform, not to be confused with MYO7A which encodes the Myosin VIIA protein utilized in our histology. FC = fold-change, p-value < 0.05.

|  |  |  |  |  |
| --- | --- | --- | --- | --- |
| Cytoskeleton | **CIB3**<br>(1.75, 4.07e-04) | Binds to TMC1 & TMC2; essential for IHC mechanotransduction | **CSRP3**<br>(-6.19, 4.83e-03) | Enables detection of mechanical stress; stabilizes actin filaments |
|  | **ESPNL**<br>(1.45, 1.03e-03) | Stereociliary protein; essential for hearing | **MYH7**<br>(-4.69, 4.08e-03) | Myosin heavy polypeptide 7; actin binding |
|  | **CDCP2**<br>(1.27, 9.63e-04) | Integral membrane protein | **XIRP1**<br>(-4.31, 4.37e-03) | Actin-binding protein homolog to stereociliary protein Xirp2 |
|  | **PTPRQ**<br>(1.09, 1.58e-03) | DFNA73; localized to stereocilia | **LRRC2**<br>(-3.92, 1.53e-03) | Encodes a member of leucine-rich repeat-containing family of proteins |
|  | **KNCN**<br>(0.84, 2.81e-03) | May be involved in stabilizing dense microtubular networks | **MYL3**<br>(-3.82, 3.16e-03) | Myosin light chain 3; atypical myosin that does not bind calcium |
|  | **ST8SIA2**<br>(0.64, 5.05e-03) | Integral membrane protein | **GDPD3**<br>(-1.36, 9.2e-11) | Glycerophosphodiester Phosphodiesterase Domain Containing 3; hydrolyzes LPA precursors |
| Transcription / Translation Factors | **LHX3**<br>(1.34, 2.52e-03) | Hair cell-specific transcription factor | **TBX6**<br>(-1.35, 1.25e-03) | T-box transcription factor involved in developmental SOX2-Notch signaling |
|  | **GFI1**<br>(0.73, 2.00e-03) | Hair cell-specific transcription factor | **GATSL2**<br>(-0.91, 1.16e-08) | Negative regulator of MTORC1 |
| Stress Response | **TPD52L1**<br>(1.02, 1.00e-05) | Positively regulates apoptosis; binds 14-3-3 | **YPEL3**<br>(-1.18, 1.41e-01) | p53 target; can trigger cell growth inhibition and senescence |
|  |  |  | **HSPA1b**<br>(-0.87, 3.52e-03) | HSP70 family member |
| Neurotransmitter Channels and Receptors | **KCNA10**<br>(1.09, 4.91e-04) | cGMP activated, voltage gated potassium channel | **GLRA1**<br>(-4.67, 2.33e-03) | Subunit of the glycine receptor |

### Loss of FOXO3 does not alter the cochlear oxidative stress response to mild noise

We utilized immunofluorescence to evaluate the oxidative stress pathway in *Foxo3^KO/KO^* and WT cochleae. No differences in HSP70 in OHCs or Deiters’ cells (DCs) were observed at baseline (Fig. S6). Immediately following noise, both genotypes similarly expressed 4-hydroxy-2-nonenal (4-HNE), in middle OHC stereocilia, indicating similar levels of lipid peroxidation (Fig. 5C, F). At baseline, JNK phosphorylation (pJNK) was highly concentrated at both IHC and OHC synapses & neurites and has lower expression in pillar and DCs (Fig. 5H, J). At 0.5 HPN, the DCs of *Foxo3^KO/KO^* and pillar cells of WT cochleae showed increased pJNK expression (Fig. 5I, K). Little to no pJNK was observed within the OHCs of either genotype. The baseline phosphorylation of cJUN (pcJUN) varied across animals but was mainly expressed in the DCs and IHC synapses (Fig. 5L, O). At 2 HPN, pcJUN was most pronounced in the DC nuclei of both genotypes (Fig. 5M, P). Low OCM+ *Foxo3^KO/KO^* OHCs contained pcJUN within their pyknotic nuclei, possibly reflecting cell stress^48^ (Fig. 5P-Q’, yellow arrows). Surprisingly, p53, which directly interacts with FOXO3^49^, showed little difference in expression at the OHCs and DCs following noise exposure (Fig. S7). These data suggest that loss of FOXO3 does not alter the cochlear oxidative stress response following noise exposure. Given these data and the presence of parthanatos, we speculate that FOXO3’s importance to OHC survivability is due to its activity prior to insult.

**Figure 5.**
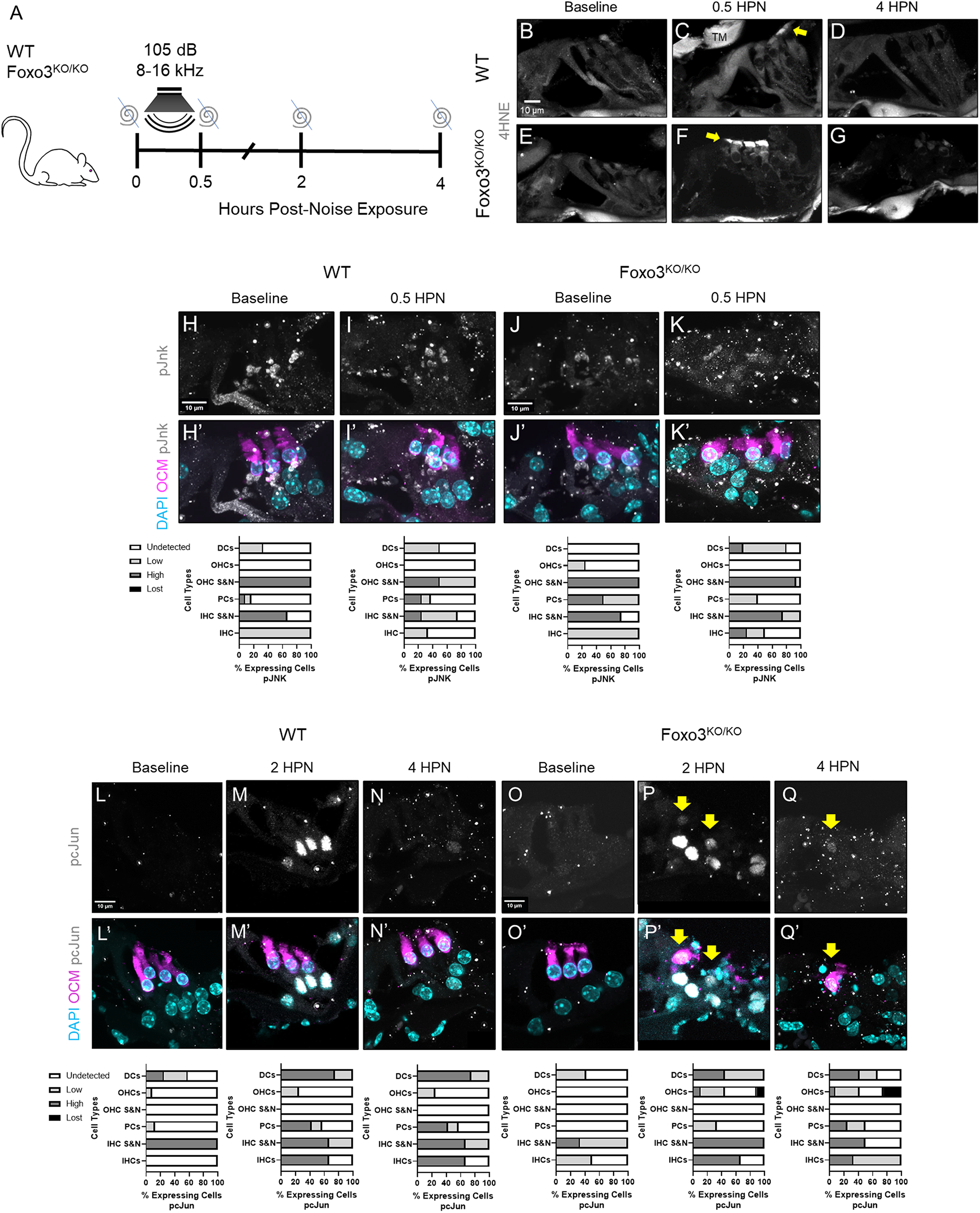
The oxidative stress response in the *Foxo3^KO/KO^* is comparable to WT littermates following noise exposure. **A)** WT and *Foxo3^KO/KO^* mice were exposed to noise and their cochleae were extracted for cryosectioning and immunohistochemistry at designated time points. **B-G)** Cochlear sections of the 24 kHz region immunostained for 4HNE (white) at baseline, 0.5 HPN, and 4 HPN. **C, F)** At 0.5 HPN, 4HNE expression increased at the OHC stereocilia in both genotypes (yellow arrows). **D, G)** 4HNE levels returned to baseline by 4 HPN. n = 3-4 per genotype/time point, 60x magnification, scale bar = 10 µm, TM = Tectorial membrane. **H-Q’)** Cochlear sections of the 24 kHz OHCs and SCs stained for DAPI (cyan) and with antibodies against OCM (magenta), pJnk (**H-K’**, white) and pcJun (**L-Q’**, white). n = 3-5 per genotype/condition, 200x magnification, scale bar = 10 µm. Corresponding % fluorescent cellular expression graphs below each set of images: undetected = white, low = light gray, high = gray, and lost cells = black; DC: Deiters’ Cells, OHCs: outer hair cells, OHC S&N: OHC synapses and neurites, PCs: pillar cells, IHC S&N: IHC synapses and neurites, and IHC: inner hair cells. (**H-K’**) Though strong at the hair cell synapses and highly variable across the organ of Corti, pJnk was absent from the OHCs in both genotypes after noise exposure. In both genotypes, pcJun expression was variable in DC nuclei at baseline (**L-L’**, **O-O’**) but strongly increased by 2 HPN (**M-M’**, **P-P’**). pcJun was present in damaged *Foxo3^KO/KO^* OHC nuclei at 2 HPN (**P, P’**), continuing through 4 HPN (**Q-Q’**) (yellow arrows).

### LPA treatment delayed but did not prevent *Foxo3^KO/KO^* OHC and hearing loss

RNA sequencing identified candidate DEGs that could contribute to increased noise sensitivity in the *Foxo3^KO/KO^* cochlea (Table 1). Actin dysregulation contributes to rapid cell death in noise damage^27^. Lysophosphatidic acid (LPA) administration can mitigate this effect by activating the ROCK2-RhoA pathway^50^. *Gdpd3* encodes an enzyme that hydrolyzes lysoglycerophospholipids to generate LPA. *Gdpd3* was downregulated at baseline in *Foxo3^KO/KO^* cochleae (Log FC = −1.36, Log CPM = 4.55, p = 9.2E-11). Decreased *Gdpd3* transcription was confirmed with RT-qPCR (fold gene expression = 0.184). Immunofluorescence showed that GDPD3 protein was expressed in neurites and supporting cells (Fig. S8). GDPD3’s proximity to OHCs makes it a candidate for intrinsic cochlear LPA production, which has not previously been identified. Unlike the RNA transcripts, we observed no difference in immunofluorescence intensity between genotypes. GDPD3 has recently been postulated to regulate FOXO3a/β-catenin binding and inhibition of the AKT/mTORC1 pathway via mediation of lysophospholipid metabolism^51^. It is possible that in the absence of FOXO3, residual GDPD3 may alter its LPA production.

We tested if cytoskeletal dysregulation contributes to rapid OHC loss in the *Foxo3^KO/KO^* by phenotypic rescue with LPA treatment. Both short and long-term analyses were performed (Fig. 6A, G). The 4 HPN cochleogram indicates that survival of OCM+ OHCs improved with LPA versus saline treatment for *Foxo3^KO/KO^* mice basal to the noise band (Fig. 6B, p = 0.0020, Mann-Whitney U). Confocal images at 24 kHz showed more OCM-negative regions with pyknotic nuclei in the saline-treated versus LPA-treated *Foxo3^KO/KO^* cochlea (Fig. 6E, F, yellow arrows). However, by 14 DPN LPA-treated and saline-treated *Foxo3^KO/KO^* mice had similar levels of OHC loss, differing from WT littermates (Fig. 6H, p < 0.0001, Kruskal-Wallis). In both groups of *Foxo3^KO/KO^* mice, ABRs following noise damage revealed significant permanent threshold shifts (Fig. 6M, p < 0.0001, two-way ANOVA). LPA treatment improved WT hearing recovery, reducing all threshold shifts back to baseline (Fig. 6M, p = 0.0495, two-way ANOVA). Identically poor DPOAEs were present in both *Foxo3^KO/KO^* groups above 16 kHz at 14 DPN (Fig. 6N, 16 kHz p < 0.0001, 24 kHz p = 0.02, 32 kHz p = 0.03, two-way ANOVA). LPA treatment improved WT OHC function versus saline-treated mice at 32 kHz (Fig. 6N, p = 0.0004, Welch’s t-test). These results suggest that a brief LPA treatment delays rapid cell death in the *Foxo3^KO/KO^* cochlea, implicating cytoskeletal dysregulation in parthanatos. LPA treatment is insufficient, however, to overcome OHC fragility or prevent hearing threshold shifts. Altogether, these results implicate FOXO3 in actin stability rather than oxidative stress response for cochlear homeostasis.

**Figure 6.**
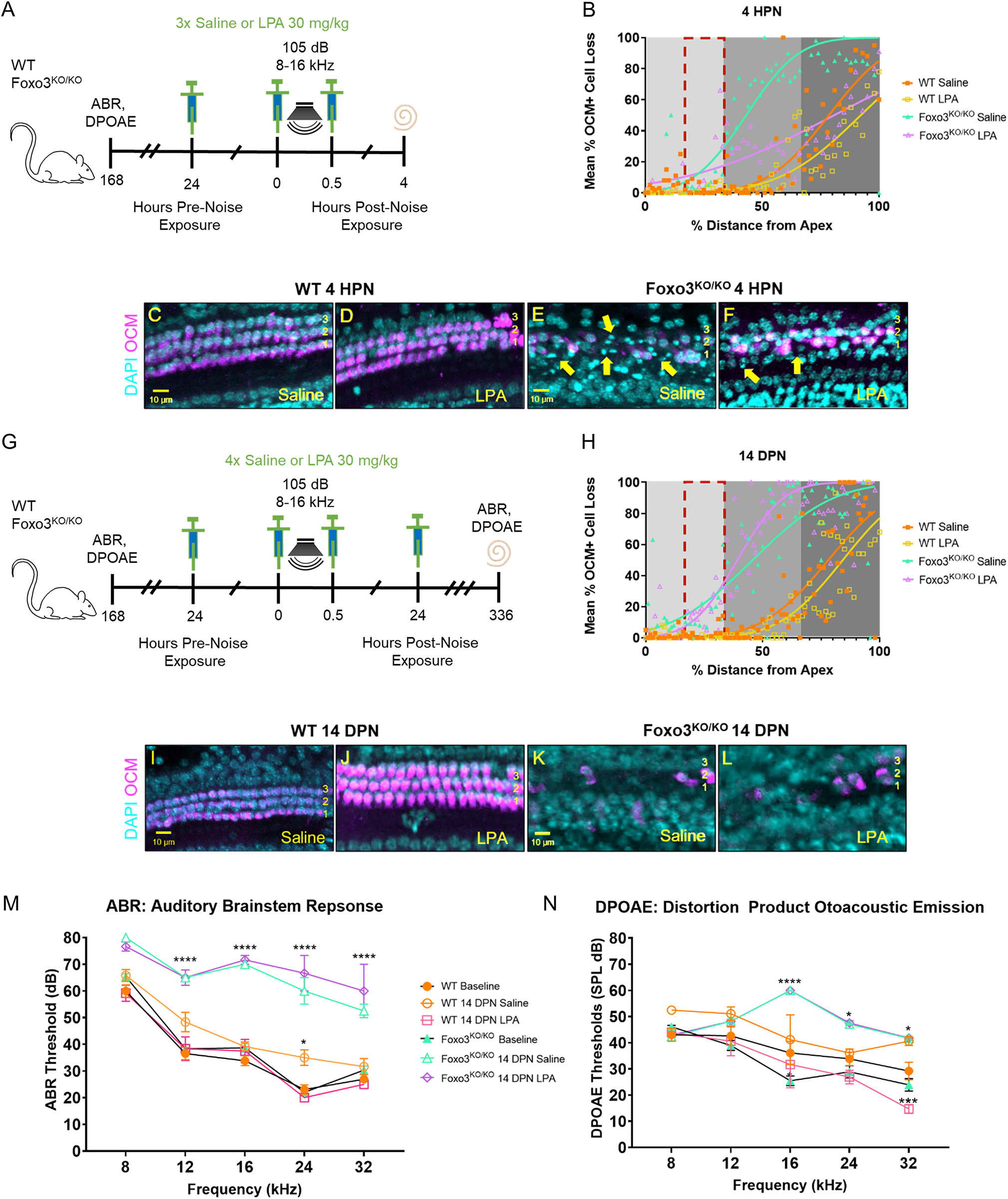
Activation of the Rho pathway with lysophosphatidic acid (LPA) delays OHC loss from noise in the *Foxo3^KO/KO^* but does not improve hearing recovery. **A, G)** All mice were tested for baseline ABRs and DPOAEs the week prior to noise exposure. **A)** One set of WT and *Foxo3^KO/KO^* mice (n = 3-5 per genotype) received three i.p. injections of LPA and were sacrificed at 4 HPN. Sham mice (n = 3-4 per genotype) received three 0.9% sterile saline injections at the same time points. Cochleae were extracted and stained for DAPI and with antibodies against OCM, and MYO7 for OHC quantification. **B)** Cochleogram at 4 HPN of % mean OCM+ cell loss (y-axis) plotted against % distance from the cochlear apex (x-axis). Each dot represents the mean number of OCM+ cells counted in 100 µM intervals along the length of cochleae (4-86 kHz) pooled (n = 3 per genotype/condition). Non-parametric interpolation lines are presented with each group: WT Saline (orange squares), WT LPA (gold squares), *Foxo3^KO/KO^* Saline (green triangles), and *Foxo3^KO/KO^* LPA (purple triangles). Cochlear tonotopic regions are presented by background color: Apex = light gray, Middle = gray, Base = dark gray. The noise band is presented as a red dotted outline. Mann-Whitney U and Kruskal-Wallis rank sum test adjusted for multiple comparisons, p < 0.05. **C-F)** Confocal images of 24 kHz OHC regions of each genotype/condition at 4 HPN immunostained with DAPI (cyan) and OCM (magenta). E, F) *Foxo3^KO/KO^* saline-treated OCM-low OHCs contained more pyknotic nuclei overall than the *Foxo3^KO/KO^* LPA-treated OHCs (yellow arrows). n = 3 per genotype/condition, 20x magnification, scale bar = 10 µm. **G)** A second set of mice (n = 3 per genotype) was injected four times with LPA and sacrificed at 336 hours (14 DPN) following hearing tests. Sham mice (n = 3 per genotype) received four saline injections at the same time points and underwent the same tests. **H)** Cochleogram at 14 DPN with the same parameters as in **B**. **I-L)** Confocal images at 14 DPN with the same parameters as in **C-F**. **K, L)** OCM+ OHC immunoreactivity in both *Foxo3^KO/KO^* groups was similar. Mean ABR (**M**) and DPOAE (**N**) thresholds (y-axis) at baseline for WT (n = 16, orange circles) and *Foxo3^KO/KO^* (n = 14, green triangles) groups, and at 14 DPN for WT Saline (n = 3, orange open circles), WT LPA (n = 3, pink open squares), *Foxo3^KO/KO^* Saline (n = 3, green open triangles), and *Foxo3^KO/KO^* LPA (n = 3, purple open diamonds) groups are plotted along five frequencies tested (x-axis). +/- s.e.m., two-way ANOVA with Tukey’s test for multiple comparisons, Welch’s t-test, alpha = 0.05, *p < 0.05, **p < 0.01, ***p < 0.001, ****p < 0.0001.

## Discussion

Irreversible loss and damage to sensory cells contributes to poor hearing outcomes in NIHL. Genetic variants in transcription factors necessary for survival and death can confer greater susceptibility to developing this disorder, including *FOXO3*^52,53^. To better understand how FOXO3 promotes cochlear homeostasis, we exposed *Foxo3^KO/KO^* mice, which lack FOXO3 in all cells throughout their lives, to mild noise. We determined the time course and cell death pathway for *Foxo3^KO/KO^* OHCs. We evaluated the effects of noise exposure on RNA expression and several damage response pathways. Finally, we tested if activation of the ROCK2-RhoA pathway could mitigate the effects of noise on *Foxo3^KO/KO^* OHCs^50^. FOXO3 is expressed in both OHCs and supporting cells and may be playing different roles in each^22^. Due to the rapidity with which OHCs are lost basal to the noise band, FOXO3 could potentially mediate expression of factors important to preserving OHCs, preventing parthanatos, and possibly enhancing cytoskeletal resilience. Contrary to our original hypothesis, we conclude that FOXO3 is not preserving the cochlea via oxidative stress regulation but rather via its transcriptional activity during development.

We show a correlation of early and prolonged OCM modulation with cell injury, possibly due to excessive calcium buffering experienced in noise exposure. OCM binds to calcium and is localized to the cuticular plate and cytoplasm of OHCs, likely to regulate their motility^38^. We found that OCM immunoreactivity normally fluctuates at baseline and in response to noise in viable OHCs (Fig. 1, S3, 2, S4). However, poor outcomes for OHCs harboring low OCM levels through 4 HPN were characteristic of the *Foxo3^KO/KO^* (Fig. 1, S3, 2, S4, 6). While not essential to cochlear development, *Ocm^KO/KO^* mice develop progressive hearing loss^39^. When modeled *in vitro*, OCM did not appear to have a direct impact on actin polymerization^54^. There have been no studies to confirm whether OCM protects against oxidative stress and whether it is responsible for multiple functions within OHCs^38^. The majority of OHCs containing fragmented nuclei expressed low OCM levels, making its loss a marker of permanent damage (Fig. 2, S4, 6).

Due to the rapidity of their demise, we consider *Foxo3^KO/KO^* OHCs primed to die. Several cell death pathways have been examined in NIHL models with apoptosis and necrosis the most prominent^44,55^. Both intrinsic and extrinsic apoptotic pathways contain various players whose actions eventually converge with caspase activation^56^. These cells are characterized by shrunken volumes and pyknotic nuclei with surface blebs, in contrast to necrotic (oncotic) cells with swollen features^31^. We saw no evidence of caspase-dependent apoptosis nor necrosis post-noise exposure (Fig. 3). We did find similarities to the “3^rd^ death pathway” as described by Bohne et al., with cells having a persistent apical region, debris arranged in a cylindrical cell-like shape with a deficient plasma membrane, and a normal sized nucleus that may later enlarge and fragment^31^ (Fig. 2, S4). Other causes of death for sensory cells have been proposed, including AIF-mediated parthanatos^45^. AIFM1 translocated to the nucleus of *Foxo3^KO/KO^* OHCs after noise exposure (Fig. 3), as previously described for guinea pig OHCs after severe, repeated acoustic insults^43^. Understanding the molecular basis for OHC fragility will lead to new models of cochlear homeostasis.

FOXO3 has previously been reported as a master regulator of oxidative stress elements in various tissues^6,7,12,57,58^. Catalase and MnSOD are two examples of reactive oxygen species-detoxifying enzymes that are decreased in haematopoietic stem cells deficient for FOXO isoforms^59^. We analyzed control cochleae for initial signs of oxidative imbalance in the *Foxo3^KO/KO^*. There was no indication of elevated lipid peroxidation, increased HSP70 accumulation in supporting cells, nor significant differential expression of stress-related genes in the *Foxo3^KO/KO^* cochlea at baseline (Fig. S6, 4, 5). The activation of the oxidative stress pathway was observed most clearly with 4HNE present in the apex of OHCs at 0.5 HPN and around 2 HPN with elevated pcJUN in supporting cell nuclei and expression in OCM-low OHCs in the *Foxo3^KO/KO^* (Fig. 5). Surprisingly, the time course and cellular expression pattern of these factors were similar between genotypes. Immediate cell damage suggests that transcription of protective elements is more crucial at baseline than the transcription of stress responders, which may be a slower process. For example, we did not see a significant downregulation in Bim at 4 HPN (Log FC = 0.10, Log CPM = 4.22, p = 0.547) which is a direct FOXO3 transcriptional target that can activate apoptosis^60^. A limitation of our RNA sequencing analysis is that we used whole cochlea lysates to analyze genetic changes and did not include the 0.5 HPN time point. FOXO3 is normally found in several cell types and its activity may be spatiotemporal. Conditional *Foxo3* knock-out lines or single cell sequencing may provide us with improved control over FOXO3 expression and cell-intrinsic versus extrinsic activity.

FOXO3 may influence OHC viability via extrinsic effects from its presence in supporting cells. We can draw a comparison to HSP70 which can be secreted immediately after noise or ototoxic insult via exosomes from supporting cells to hair cells for their protection^61–66^. Seeing that *Gdpd3*, one of the only genes downregulated in the *Foxo3^KO/KO^* cochlea at baseline, is strongly expressed in the supporting cells (Fig. S8), we speculate that FOXO3 may have a role in its transcription or activation. The fact that LPA was found to delay OCM changes but not completely rescue OCM numbers supports the idea that the LPA-activated ROCK2-RhoA stabilization of actin filaments can mitigate some of the damage in our noise exposure (Fig. 6). However, the differential expression of cytoskeletal genes correlated with *Foxo3* deletion (Tables 1, S3) may also decrease overall actin stability in the OHCs and/or supporting cells. Further investigation into cytoskeletal changes including measuring F/G actin ratios will help clarify these observations. To determine if GDPD3 can contribute to NIHL, we would like to evaluate *Gdpd3^KO/KO^* mice for cochlear LPA levels and hearing thresholds after noise exposure. Recapitulation of the *Foxo3^KO/KO^* phenotype would aid in our understanding of the overall mechanism of action for maintaining OHC viability.

In summary, this study provides valuable insight into the mechanism underlying OHC loss in the *Foxo3^KO/KO^* cochlea after mild noise exposure. Future studies will focus on clarifying FOXO3’s cell-intrinsic versus extrinsic functions and its possible influence on cytoskeletal integrity.

## Materials and Methods

### Mice

Mice were housed on a 12-hour light/dark cycle and received chow and water *ad libitum*. Once pups were found, their appearance was compared with the Jackson Labs pups’ age appearance chart and their birth date was designated postnatal day (P) 0. Mice were weaned between P21-28 and no more than five adults were housed in the same cage. They were provided ample nesting materials and small houses within their home cages. Using the NIOSH Sound Level Meter app (Centers for Disease Control, Washington, DC, USA), ambient noise in the mouse colony room was estimated at approximately 70 dB with interior cage levels centered around 58 dB. Cages to be used for experiments were maintained in the center of the holding rack to avoid excess noise.

Mice were genotyped and then randomly assigned to their conditions using a deck of cards. For in-house genotyping, DNA was obtained from 2-mm tail samples digested overnight in Proteinase K (IBI Scientific, Dubuque, IA, USA) solution at 65°C followed by phenol/chloroform extraction. IProof Taq (BioRad, Hercules, CA, USA) was used in conjunction with a published protocol and primer sequences^67^. For outsourced genotyping, DNA obtained from 2-mm tail samples were processed and verified by Transnetyx Automated PCR Genotyping Services (Transnetyx, Inc. Cordova, TN, USA).

Heterozygous/homozygous FoxO3 (FoxO3^+/2^) mice (FVB;129S6-Foxo3tm1.1Rdp; RRID:MGI:2668335) were obtained from the Mutant Mouse Regional Resource Center, at the University of California at Davis (Davis, CA, USA; stock no. 016132-UCD)^68^. Originally made on 129 Sv and bred three times to FVB/n, we bred the line a fourth time to FVB/nJ (RRID:IMSR JAX:001800) to obtain heterozygote mutant mice. Heterozygotes with different parents were bred together to obtain both knockout and *Foxo3^+/+^* littermates. *Foxo3^KO/KO^, Foxo3^+/KO^,* and *Foxo3^+/+^* littermates, both male and female, between the ages of P60-90 were used in this study. *Foxo3^+/+^* littermates are denoted as wild-type (WT) mice.

### Noise Exposure

Mice were exposed to an 8-16 kHz octave noise band at 105 dB for 30 minutes. This exposure has previously been shown to induce temporary threshold shifts in WT mice and loss of hearing in *Foxo3^KO/KO^* mice after 14 days when measured at five frequencies^23^. Awake P60-90 mice were placed in individual triangular wire mesh cages, 12 cm x 5 cm x 5 cm, in an asymmetric plywood box with a JBL2250HJ compression speaker and JBL2382A biradial horn mounted above. This apparatus was contained within a sound booth. The speaker was driven by a TDT RX6 multifunction processor and dedicated attenuator (Tucker Davis Technologies, Alachua, FL, USA). It was controlled with TDT RPvdsEx sound processing software (Tucker Davis Technologies). All noise exposure equipment was calibrated prior to use via the Quest Technologies portable sound level meter, Model 1900 (TSI Incorporated, Shoreview, MN, USA). The sound level was checked with an iPhone using the NIOSH SLM app and the iPhone’s internal microphone prior to each exposure. The iPhone was previously calibrated with the SoundMeter app software and a solid state 94 dB source (provided with the Quest sound meter) in a sound booth. Within the sound exposure box, cages were placed in three specific locations at the back of the box where sound levels were highly consistent (+/− < 0.5 dB). Mice that escaped or moved their cages from the starting position were excluded. Mice were directly monitored for the first minute of exposure via the plexiglass window on the chamber. A minority of FVB/nJ mice were susceptible to audiogenic seizures^69^, normally within the first minute of exposure. These mice were immediately removed from the chamber and excluded. Due to known circadian cycle interactions with noise damage, mice were exposed to noise only between the hours of 9am and 1pm^70^. Control mice were placed in the apparatus for the same length of time with the power off. After noise exposures, mice were monitored for proper health status and closely examined up until sacrifice. Mice exhibiting signs of pain or distress were euthanized early and excluded from further analysis.

### Auditory Brainstem Response (ABR) and Distortion Product Otoacoustic Emission (DPOAE)

Initial hearing thresholds were tested between ages P50-55 with additional testing following schedules as described in the text. Testing occurred between the hours of 9am – 6pm to avoid circadian rhythm effects. Mice were anesthetized with a single intraperitoneal (i.p.) injection of ketamine (80 mg/kg animal weight) and acepromazine (3 mg/kg animal weight) diluted in a sterile saline solution, to provide approximately 45 minutes of immobility. Additional anesthetic was administered as needed. A 10B+ OAE microphone was housed in an interaural probe and coupled with the speaker outputs. The probe was placed at the opening of each mouse’s left external auditory meatus. A Smart EP Universal Smart Box (Intelligent Hearing Systems, Miami, FL, USA) with an ED1 speaker (Tucker Davis Technologies) were housed within an anechoic chamber and used for closed field auditory testing. The Quest Technologies portable sound level meter, Model 1900, was used to calibrate the apparatus less than one month prior to beginning each set of experiments.

The auditory brainstem response (ABR) consisted of 5-ms click stimuli followed by 5-ms tone pips presented at five frequencies (8, 12, 16, 24, and 32 kHz). Stimuli amplitudes decreased in 5 dB steps from 75 dB sound pressure level to 15-25 dB. The averages of 512 sweeps were recorded for each frequency and amplitude. Three sterilized fine subdermal electrodes were used to record electrical responses (Grass): one inserted at the vertex and one inserted beneath each pinna. Responses were rejected if their peak to trough amplitude was greater than 31 µV at any time between 1.3 to 12.5 msec after stimulus presentation. Well-anesthetized mice typically had a 5-30% rejection rate. ABR thresholds at each of the five frequencies were determined by the last visible trace of wave I (dB). If no waveform was observed, “80 dB” was designated as the ceiling threshold as it can be considered reflective of severe hearing loss and higher intensities might produce acoustic distortions^71,72^.

Distortion product otoacoustic emissions (DPOAEs) were measured using the amplitude of evoked otoacoustic emissions to simultaneous pure tones of frequencies f1 and f2, where f1/f2 = 1.2 and the f1 level is 10 dB above f2. Beginning with f1 at 20 dB and ending at 65 dB, 32 sweeps were made in 5 dB steps. The DPOAE threshold was calculated for 3 dB emission. As a second calibration, measurements were collected from a dead mouse and L2 amplitudes with signals above threshold were excluded.

Anesthetized mice were isolated in recovery cages until they woke up. Their arousal levels were monitored, and mice were returned to their home cages after they regained consciousness. The researchers scoring ABRs and DPOAEs were blinded to genotype, condition, and time point.

### Tissue Preparation for Immunostaining

Cochleae were dissected from euthanized mice at designated time points. The stapes was removed, and a small incision was made at the apical tips for proper fluid exchange during immersion fixation with 4% paraformaldehyde (PFA)/1x phosphate-buffered saline (PBS). Tissues were transferred to decalcifying 0.1 M ethylenediaminetetraacetic acid (EDTA) at 4°C on a rocker for three days. For cryosectioning, tissues were submerged in 30% sucrose/1x PBS overnight, embedded in optimal cutting temperature compound (OCT), and frozen with liquid nitrogen. These tissues were sectioned at 20 μm with the 12 and 24 kHz regions visible. The 12 and 24 kHz regions were selected to analyze changes within the noise band (8-16 kHz) and basal to it.

For whole mount preparations, cochleae were microdissected into three turns (apical, middle, and basal) as previously described^73^. These pieces were frequency mapped using the ImageJ 64 (NIH) plug-in from Massachusetts Eye and Ear Infirmary, and immunostained for quantification and spatiotemporal analysis. Apical IHCs have high levels of oncomodulin^74^ and can crowd areas of OCM+ OHCs in the apical hook region (< 4 kHz), making differentiation difficult. For our cochleograms, we included cell counts from 4 kHz to 86 kHz (0-100% distance from the apex)^75^.

### Primary Antibodies

Primary antibodies are listed in Supplementary Table 1. All antibodies were initially tested using antigen and non-antigen retrieval via freezing and boiling to ensure adequate cellular penetrance and stability. Non-primary and non-secondary antibody control tissues were imaged for autofluorescence levels. Due to a manufacturer’s discontinuation, we utilized two oncomodulin antibodies in this study. Both antibodies were compared with and without antigen retrieval steps on control tissues. No difference in fluorescent antigen binding was noted.

### Immunohistochemistry

For immunohistochemical staining on cryosections, sections were washed with 0.5% Tween20/Tris-buffered saline (TTBS), pH 7.4, and then blocked for 1 hour at room temperature in 5% normal donkey serum (Jackson ImmunoResearch, West Grove, PA, USA)/0.5% TTBS. For anti-HSP70 immunostaining, antigen retrieval was performed prior to these steps by boiling in 10 mM citric acid, pH 6, for 15 min at 20% microwave power. Antibody incubations were performed overnight at 4°C in block solution. A complete list of primary antibodies is provided in Supplementary Table 1. Sections were washed in 0.5% TTBS at room temperature prior to secondary antibody incubation overnight at 4°C in the dark. Alexafluor-conjugated secondary antibodies included 488, 594, and 647 (Thermo Fisher Sci, Waltham, MA, USA) and were diluted at 1:500 each. 4’6-diamidino-2-phenylindole (DAPI) diluted at 1:10000 was added during the secondary antibody incubation. Sections were washed in 0.5% TTBS and mounted in ProLong Gold Antifade (Thermo Fisher Sci) on frosted plus microscope slides (Thermo Fisher Sci).

For whole mount preparations, tissues were collected in 300 μL dPBS in Eppendorf tubes, flash frozen with liquid nitrogen, thawed on ice, and transferred to a 40-well plate. Tissues were washed in 0.5% TTBS, blocked for 1 hour at room temperature in 5% normal donkey serum/0.5% TTBS, and incubated in 200 μL of primary antibody overnight at 4°C. Tissues were washed in 0.5% TTBS at room temperature and incubated in 200 uL of secondary antibody solution overnight in the dark at 4°C. Tissues were washed with 0.5% TTBS prior to mounting in ProLong Gold Antifade between two 50 mm coverslips (Thermo Fisher Sci).

### Confocal & Multiphoton Microscopy and Image Processing for Figures

Sections were imaged using an Olympus FV1000 (Olympus, Tokyo, Japan) and ZEISS LSM 510 Meta (Carl Zeiss AG, Oberkochen, Germany) laser scanning confocal microscopes. Whole mounts were imaged using the confocal microscopes and Olympus Fluoview FVMPE-RS multiphoton imaging system. All images presented were collected as z-stack projections using one assigned microscopy device for appropriate comparison. Stitching was performed with Fluoview for multiphoton images or XuvStitch (XuvTools) for confocal images. Composite images of whole mounts were assembled in GIMP 2.10.12 using the layering function to match the frequency regions previously mapped in ImageJ 64 (NIH) after microdissection. For cochleograms, 100 μm bins were measured along the length of the mapped cochlea on the rows of pillar cells. Oncomodulin (OCM), Myosin 7a (MYO7), and Cytochrome-C were used to denote the presence of OHCs and IHCs. Any areas of the cochlea with post-mortem damage to the sensory epithelium were omitted from the analysis. These regions of the scatter plots are empty. In regions with intact sensory epithelium and high OHC loss, an average OHC count determined from the cochlea’s low-frequency regions was used as the denominator^75^. For analysis of the oxidative stress response, normalized fluorescence for sections was performed in ImageJ and relative expression levels were categorized as undetected, low, high, and cells lost. Standardization of counts involved analyzing the same number of cells per section (1 IHC, 1 cluster of IHC synapses and neurites, 3 OHCs, 3 clusters of OHC synapses and neurites, 2 pillar cells, and 3 Deiters’ cells) before calculating total percentages of those surveyed. For analysis of Hsp70 levels in OHCs and DCs, images were opened in ImageJ, LUT was set to 0-75, and the corrected total cell fluorescence (CTCF) was calculated using the formula CTCF = Integrated Intensity – (Area of selected cell x Mean fluorescence of background readings).

### 3-D Reconstructions of Outer Hair Cells and Nuclei

To assess the structural integrity of OHCs and their nuclei following noise damage, 12 and 24 kHz regions from the previously mapped cochleograms were imaged at 150x on the Olympus FV1000. These frequencies were selected to match the results in cryosection. All regions were standardized in the x, y, and z planes to include the expected three rows of OHCs. Images were converted (.oif to .ims) and imported into Imaris 9.3 Image Visualization & Analysis Software (Oxford Instruments, Abingdon, Oxfordshire, UK) for 3-D reconstruction. All sensory epithelial sections were oriented in the same direction and any remaining tectorial membrane was cropped out of the visualization frame. DAPI (nuclei), oncomodulin (OHCs), and Cytochrome-C (OHCs) isosurfaces were appropriately thresholded and constructed. Despite individual isolation attempts, thresholding was incapable of separating all supporting cell nuclei proximate to those of the OHCs. The resulting images have supporting cell nuclei designated as “SC” unless otherwise stated in the figure legend. Total OHC numbers were graphed for cells expressing each fluorescent marker.

### RNA Sequencing

At P60, WT and *Foxo3^KO/KO^* mice were euthanized under control conditions or at 4 hours post-noise (HPN) (n = 6 per genotype/condition). Their cochleae were extracted and processed for RNA sequencing. RNA was isolated and purified using the RNeasy Plus Mini Kit (QIAGEN, Hilden, Germany). A 2100 Bioanalyzer (Agilent, Santa Clara, CA) was used to assess quality. The minimum RNA integrity number (RIN) for the group was 8.3 and the average was 9.1. Samples were processed at the University of Geneva, Switzerland. RNA was sequenced on an Illumina HiSeq4000 (Illumina, San Diego, CA, USA). The reads were mapped with TopHat v.2 software (CCB, Johns Hopkins University, Baltimore, MD, USA) to the UCSC mm10 mouse reference (Genomics Institute, University of California Santa Cruz, Santa Cruz, CA, USA). Biological quality control and summarization were performed with RSeQC-2.3.3 and PicardTools1.92. The table of counts was prepared with HTSeq v0.6p1. Differential expression analysis used the statistical analysis R/Bioconductor package EdgeR v. 3.10.5. Counts were normalized according to the library size and filtered. Gene expression clustering was performed using the R/Bioconductor Mfuzz v.2.28 package. These genes were composed in Supplementary Table 3 and their listed functions were adapted from the GeneCards human gene database (Weizman Institute of Science, LifeMap Sciences). Gene ontology analyses on additional gene sets were performed in Ingenuity Pathway Analysis (QIAGEN) and GSEA. Genes expressed in OHCs were identified through comparison with the gEAR portal (He dataset).

### Real Time Quantitative PCR (RT-qPCR)

At P60, WT and *Foxo3^KO/KO^* mice were euthanized under control conditions or at 4 HPN (n = 4-5 per genotype/condition). Cochleae were isolated and total RNA was extracted using the RNeasy Plus Mini Kit (QIAGEN). Total RNA concentration and purity was determined with the NanoDrop 1000 spectrophotometer (NanoDrop, Wilmington, DE, USA). A qScript cDNA Synthesis kit (QuantaBio, Beverly, MA, USA) and T100 Thermal Cycler (Bio-Rad Laboratories, Inc., Hercules, CA, USA) were used to generate the cDNA. RT-qPCR was performed with SYBR Green PCR mix on the Bio-Rad CFX Connect Real-Time PCR System (Bio-Rad). The reaction protocol used was 95 °C for 30 s, followed by 50 cycles of 95 °C for 15 s, 60 °C for 60 s, and 4 °C to hold. Comparative ΔΔCt was performed to determine relative mRNA expression, where relative abundance of each gene was first internally normalized to the geometric mean Ct for the reference genes Hprt and GusB^76^. The primers are listed in Supplementary Table 2.

### Administration of LPA to mice

The selected lysophosphatidic acid (LPA) concentration was considered sufficient for OHC preservation following noise exposure based on previous findings in adult CBA/J mice^50^. LPA (30 mg/kg, Oleoyl-L-α-lysophosphatidic acid sodium salt, Sigma L7260-25MG) was prepared as 0.05 mg/µL stock with CMF-PBS and diluted in 0.9% sterile saline at a final 5 mg/mL concentration. One set of WT and *Foxo3^KO/KO^* mice (n = 3-5 per genotype) received a total of three i.p. injections of LPA and were sacrificed at 4 HPN. LPA was injected 24 hours and immediately prior to noise exposure, and immediately after noise exposure. Sham mice (n = 3-4 per genotype) received three 0.9% sterile saline injections at the same time points. A second set of mice (n = 3 per genotype) was injected four times with LPA and sacrificed at 14 DPN following hearing tests. LPA was administered 24 hours and immediately prior to noise, and immediately and 24 hours after noise exposure. Sham mice (n = 3 per genotype) received four saline injections at the same time points.

### Statistical Analyses

Sample sizes were calculated for sufficient power analysis and at least three cochleae of each genotype were analyzed for all experimental conditions. The number of replicates is indicated within each figure legend. The researcher was blinded as to the genotype and condition for the quantification of outer hair cell counts, immunohistochemistry fluorescence labeling, and hearing test analyses. Unpaired and paired student’s t-tests or Mann-Whitney U rank sum tests were used to compare differences across two groups. Two-way ANOVA or Kruskal-Wallis rank sum tests were used to compare differences across more than two groups. Bonferroni, Tukey’s, Šidák’s, and Welch’s post-tests were used for adjustments. A p value < 0.05 was considered statistically significant. Statistics were calculated in GraphPad Prism 9.0.0.

## Acknowledgements

We thank the University of Geneva Genomics Core for our RNA sequencing; Dr. Helene McMurray for her assistance in GO analysis; Dr. Paivi Jordan for microscopy training; technical director Dr. V. Kaye Thomas and Julie Zhang of the URMC Center for Advanced Light Microscopy and Nanoscopy, for their assistance in confocal imaging; technical director Dr. Yurong Gao of the URMC Multiphoton and Analytical Imaging Center for help with multiphoton imaging; Drs. Richard Libby, Lin Gan, Ruchira Singh, and Ian Dickerson for equipment access; Drs. Joseph C. Holt, Robert Freeman, Dirk Bohmann, and Amy Kiernan for advice on experimental planning; and Dr. Dorota Piekna-Przybylska, Dr. Hitomi Sakano, and Daxiang Na for editing.

## Conflict of Interest

The authors declare no known conflicts of interest nor competing financial interests associated with this publication.

## Author Contributions

H.J.B. and P.M.W. performed study concept, design, and analysis of RNA sequencing results; H.J.B. developed the methodology, provided acquisition, analysis and interpretation of data, statistical analysis, and writing, review, and revision of the paper; F.G. developed the methodology and processed the tissue for the RNA sequencing experiments; J.Z., S.J., and P.M.W. scored the hearing tests; J.Z. provided valuable feedback and assistance on performing experiments; P.M.W. provided technical and material support. All authors read and approved the final paper.

## Ethics Approval

All experiments were performed under protocol number 2010-011: PI Patricia White, in compliance with the U.S. Department of Health and Human Services and were reviewed by the University of Rochester’s Committee on Animal Resources.

## Funding

This work was funded by the National Institute of Health R01 DC014261.

## Data Availability

The datasets used and/or analyzed during the current study are available from the corresponding author on request. The datasets generated and/or analyzed during the current study are available in the University of Rochester repository.

## Notes

### Competing Interest Statement

The authors have declared no competing interest.

