## Supplemental tables and figures for "Primed to Die: An Investigation of the Genetic Mechanisms Underlying Noise-Induced Hearing Loss and Cochlear Damage in Homozygous Foxo3-knockout Mice"

| Name | Primary Antibody (catalog no.) | Concentration | Company |
|---|---|---|---|
| Oncomodulin (OCM) | discontinued | 1:500 | Santa Cruz |
| | (PA5-47832) | 1:500 | Thermo Fisher Sci |
| MYO7 | 25834) | 1:500 | Santa Cruz |
| Cytochrome-C | c (556433) | 1:500 | BD Biosciences |
| CASP3 | caspase-3 (9664) | 1:1000 | Cell Signaling |
| AIFM1 | inducing factor 1-mitochondrial (NBP2-33932) | 1:1000 | Novus Biologicals |
| pJnk | Monoclonal rabbit anti-pJNK (4668) | 1:250 | Cell Signaling |
| pcJun | Jun (3270) | 1:250 | Cell Signaling |
| p53 | AP) | 1:500 | Proteintech |
| 4HNE | Monoclonal mouse anti-4-Hydroxynonenal (MAB3249) | 1:500 | R&D Systems |
| Hsp70 | Polyclonal rabbit anti-HSP70 (4872) | 1:200 | Cell Signaling |
| GDPD3 | 59581) | 1:50 | Thermo Fisher Sci |

| Name | Forward Sequence | Reverse Sequence |
| --- | --- | --- |
| Hprt | 5’-TCAGTCAACGGGGGACATAAA-3’ | 5’-GGGGCTGTACTGCTTAACCAG-3’ |
| GusB | 5’-GGCTGGTGACCTACTGGATTT-3’ | 5’-GGCACTGGGAACCTGAAGT-3’ |
| Junb | 5’-TCACGACGACTCTTACGCAG-3’ | 5’-CCTTGAGACCCCGATAGGGA-3’ |
| Gdpd3 | 5’-CTCTCCTGTACTTTGTTCTGCC-3’ | 5’-CCAGGCGGATAGGGAAGAC-3’ |

**Figure Legend S1.**

**Supplementary Figure 1. Anatomy of the organ of Corti and mouse cochlear tonotopy. A)** Schematic cross-section of the main cell populations within the organ of Corti. Sensory cells detect acoustic stimuli using stereocilia that bend due to tectorial membrane displacement and basilar membrane movement. Outer Hair Cells (OHCs, red) are arranged in three rows and act as amplifiers for acoustic signal transduction by one row of Inner Hair Cells (IHC, yellow). Efferent nerves, synapsing onto the OHCs and IHCs, and afferent nerves, synapsing onto the IHCs, bundle together to form the Auditory Nerve (gray). Several supporting cells aid in the homeostatic maintenance of the sensory cells surrounding the tunnel of Corti (ToC); these cells include the Border Cells (Borders), Inner Phalangeal Cell (IPhC), Pillar Cells (PCs), Deiters' Cells (DCs), Hensen's Cells (HCs), Boettcher Cells (BCs), and Claudius Cells (CCs) (cyan). This organization continues along the entire length of the cochlea. **B)** Diagram of estimated hearing frequencies in mouse cochlea. The mouse cochlea is arranged tonotopically and can detect frequencies up to 100 kHz. Low frequencies range from 1-16 kHz along the apex, middle frequencies from 17-32 kHz, and high frequencies from 33-100 kHz in the base.

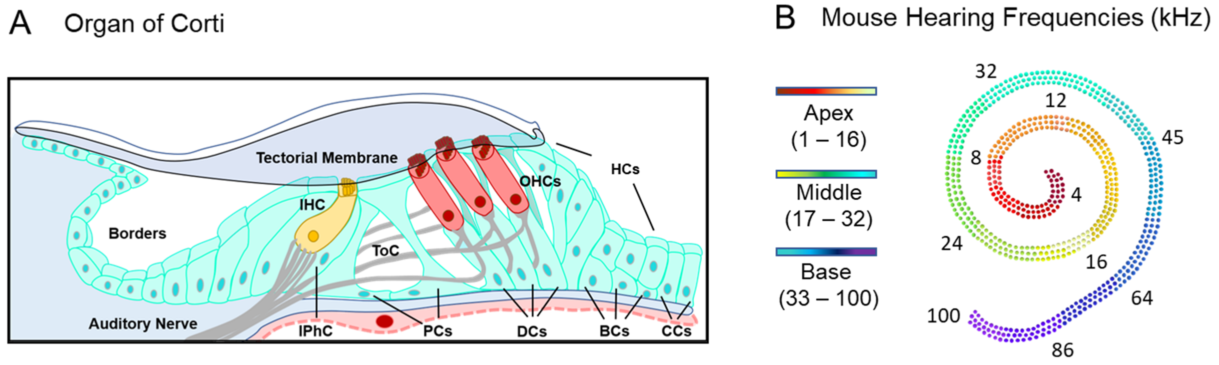
A
Organ of Corti
Tectorial Membrane
HCs
OHCs
IHC
Borders
ToC
Auditory Nerve
IPhC
PCs
DCs
BCs
CCs
B
Mouse Hearing Frequencies (kHz)
Apex
(1 – 16)
Middle
(17 – 32)
Base
(33 – 100)
32
12
45
8
4
24
16
64
100
86

**Figure Legend S2.**

**Supplementary Figure 2.** ***Foxo3*$^{+/KO}$ mice experience temporary threshold shifts after mild noise exposure. A)** *Foxo3*$^{+/KO}$ mice were subjected to an 8-16 kHz noise band at 105 dB for 30 min. Hearing tests were performed at baseline (-5 DPN), 1 DPN, and 14 DPN. **B)** ABR thresholds (y-axis) for individual *Foxo3*$^{+/KO}$ mice are plotted along five frequencies tested (x-axis). Means are represented as bars for baseline (black circles, pink bars), 1 DPN (gray squares, cyan bars), and 14 DPN (light gray triangles, green bars). Significant threshold shifts were observed at 1 DPN at 16 kHz (**p = 0.0032), 24 kHz (****p < 0.0001) and 32 kHz (****p < 0.0001). ABR thresholds at these frequencies returned to baseline by 14 DPN. n = 6-8, two-way ANOVA with Bonferroni's multiple comparisons test, alpha = 0.05, *p < 0.05, **p < 0.01, ***p < 0.001, ****p < 0.0001. Females had better thresholds at 12 kHz (*p = 0.0266) but all other comparisons indicated similar threshold shifts and recovery between the sexes (2-3 males, 4-5 females, two-way ANOVA with Šidák's multiple comparisons test, alpha = 0.05, *p < 0.05). **C)** DPOAE thresholds with the same parameters as in **B**. Significant threshold shifts were observed at 1 DPN at 16 kHz (****p < 0.0001), 24 kHz (***p = 0.0003) and 32 kHz (****p < 0.0001). By 14 DPN at all frequencies, no significant differences were detected compared to baseline. n = 6-8, two-way ANOVA with Bonferroni's multiple comparisons test, alpha = 0.05, *p < 0.05, **p < 0.01, ***p < 0.001, ****p < 0.0001. Females had better 1 DPN thresholds at 16 kHz (*p = 0.0375) but no other significant differences were observed between the sexes (2-3 males, 4-5 females, two-way ANOVA with Šidák's multiple comparisons test, alpha = 0.05, *p < 0.05).

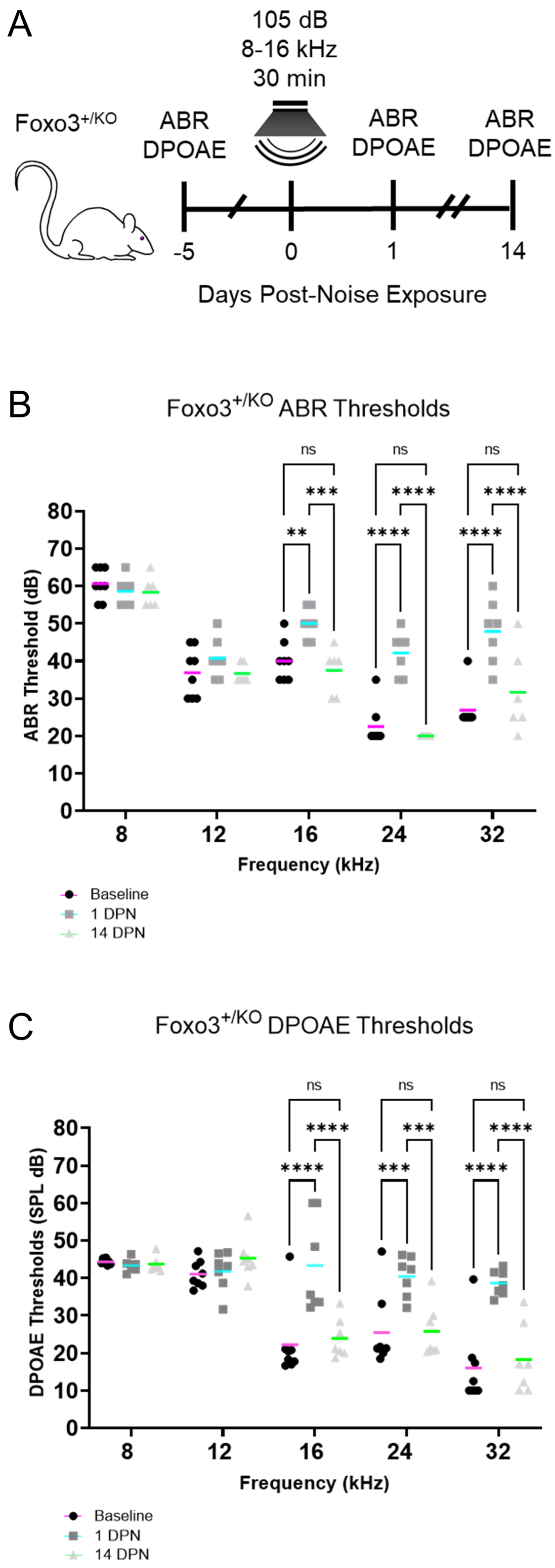

A
105 dB
8-16 kHz
30 min
Foxo3+/KO
ABR
DPOAE
ABR
DPOAE
ABR
DPOAE
-5
0
1
14
Days Post-Noise Exposure
B
Foxo3+/KO ABR Thresholds
ABR Threshold (dB)
Frequency (kHz)
Baseline
1 DPN
14 DPN
C
Foxo3+/KO DPOAE Thresholds
DPOAE Thresholds (SPL dB)
Frequency (kHz)
Baseline
1 DPN
14 DPN

**Figure Legend S3.**

**Supplementary Figure 3. At 24 kHz, *Foxo3*$^{+/KO}$ mice show mild loss of OCM immunoreactivity following noise.** After immunostaining in whole mount, the 24 kHz cochlear region was mapped, imaged using confocal microscopy at 150x magnification, and rendered in Imaris (n = 3 cochleae per genotype/condition). **A-D)** The rendered images of OHCs colored cyan for DAPI+ cell nuclei. **A'-D')** Renderings of DAPI+/OCM+ OHCs with OCM in magenta. **A"-D")** Renderings of DAPI+/OCM+/CytC+ cells with CytC in gray. n = 3 cochleae, scale bar = 5 µm, SC = supporting cell nuclei, TM = tectorial membrane. **B-B")** At 0.5 HPN, a minority of *Foxo3*$^{+//KO}$ OHCs expressed less OCM immunoreactivity, mainly in rows 2 and 3 (yellow arrows). **C-C")** Only one OCM-/CytC+ OHC remained in row 3 by 4 HPN (yellow arrows). **D-D")** OCM expression and OHC numbers were comparable to controls.

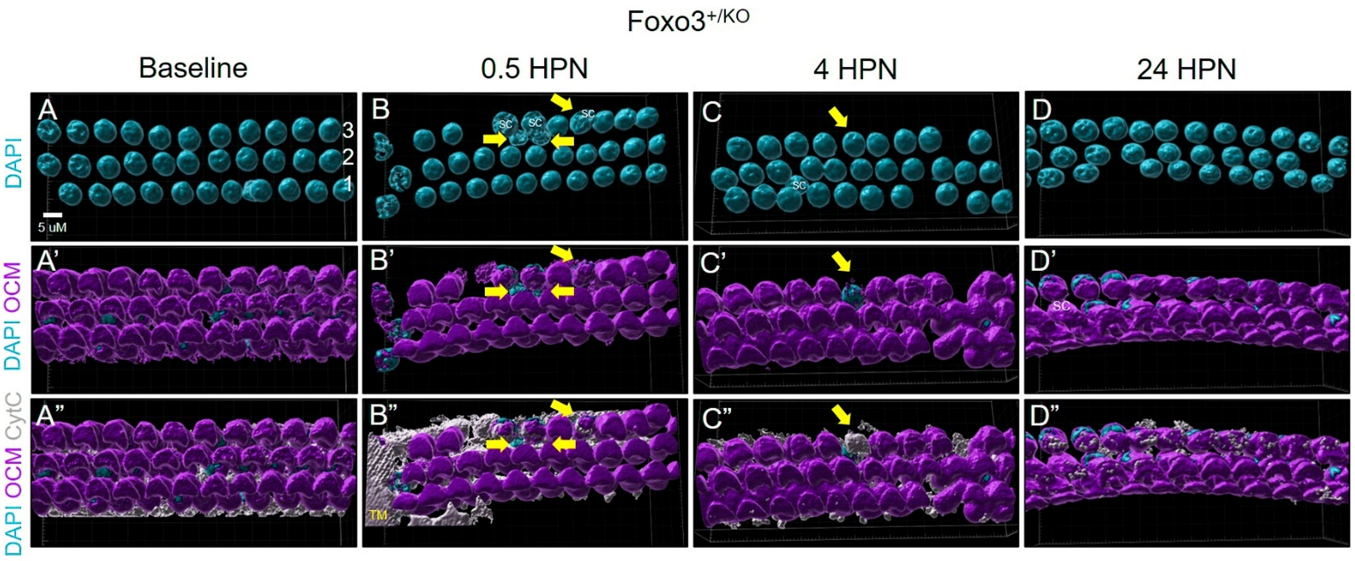
Foxo3+/KO
Baseline
0.5 HPN
4 HPN
24 HPN
DAPI
DAPI OCM
DAPI OCM CytC
A
A'
A"
B
B'
B"
C
C'
C"
D
D'
D"
3
2
1
5 uM
SC
TM

**Figure Legend S4.**

**Supplementary Figure 4. Within the noise band, *Foxo3*$^{KO/KO}$ OHCs undergo OCM modulation but little cellular loss by 24 HPN.** Following whole mount histology, the 12 kHz cochlear region was mapped, imaged with confocal microscopy at 150x magnification, and rendered in Imaris (n = 3 cochleae per genotype/condition). **A-L)** The rendered images of OHCs DAPI+ cell nuclei in cyan. **A'-L')** Renderings of DAPI+/OCM+ OHCs with OCM in magenta. **A"-L")** Renderings of DAPI+/OCM+/CytC+ cells with CytC in gray. Scale bar = 5 µm, TM = remnants of tectorial membrane, SC = supporting cell nuclei. **A-D")** WT OHCs, **E-H")** *Foxo3*$^{+/KO}$ OHCs, **I-L")** *Foxo3*$^{KO/KO}$ OHCs. **A-D")** WT OHCs showed no differences in OCM or CytC expression from baseline through 24 HPN. **E"-G")** *Foxo3*$^{+/KO}$ OHCs did not show strong changes in OCM immunoreactivity until 4 HPN (yellow arrows). **H-H"**) By 24 HPN, OCM expression was comparable to baseline levels in *Foxo3*$^{+/KO}$ OHCs. **J-J")** At 0.5 HPN, the *Foxo3*$^{KO/KO}$ OHCs became disordered and several lost OCM immunoreactivity (yellow arrows). **K-K")** Pyknotic nuclei were present in OHCs with low OCM levels by 4 HPN (yellow arrow). **L-L")** OHCs without OCM showed slightly enlarged nuclei and thin cell bodies containing CytC (yellow arrows). **M-O)** OHC counts for all three genotypes expressing specific fluorophores (y-axis) were graphed versus the noise exposure timeline (x-axis). WT (black circles, orange mean bar), *Foxo3*$^{+/KO}$ (gray squares, pink mean bar), and *Foxo3*$^{KO/KO}$ (light gray triangles, green mean bar) total cell counts per 1024 x 400 x 20-pixel selection. **M)** Numbers of DAPI+ OHC nuclei did not significantly differ between genotypes following noise. **N)** Counts of OHCs as calculated by totaling OCM+/CytC+ cells showed significant OCM modulation in *Foxo3*$^{KO/KO}$ OHCs through 4 HPN. **O)** OHCs only expressing CytC increased in the *Foxo3*$^{KO/KO}$ mice at 0.5 HPN but were comparable to baseline levels by 24 HPN. n = 3 per genotype/condition, two-way ANOVA with Tukey's test for multiple comparisons, alpha = 0.05, *p < 0.05, **p < 0.01, ***p < 0.001, ****p < 0.0001.

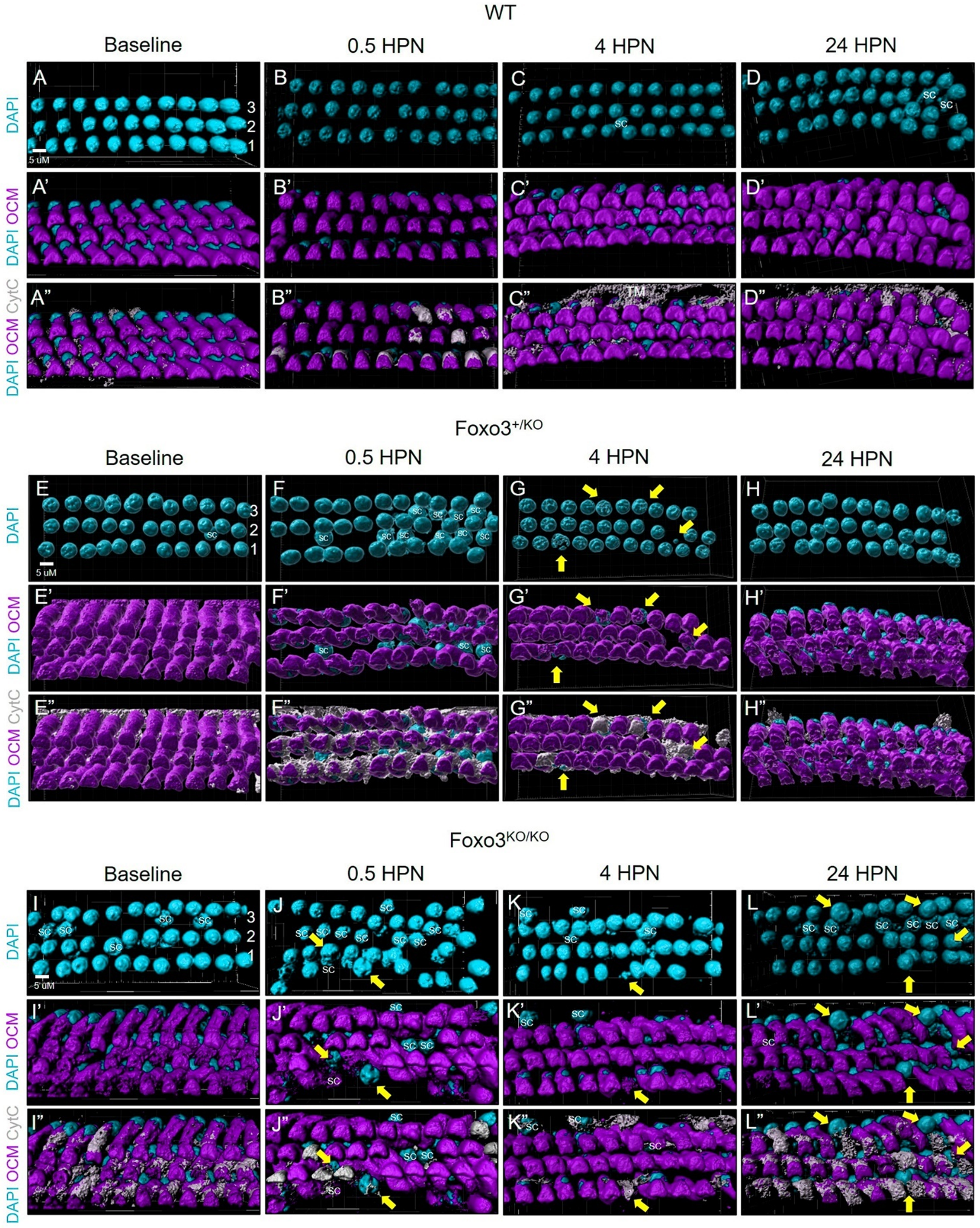

WT
Baseline
0.5 HPN
4 HPN
24 HPN
DAPI
DAPI OCM
DAPI OCM CytC
A
B
C
D
A'
B'
C'
D'
A"
B"
C"
D"
3
2
1
5 uM
sc
TM
Foxo3+/KO
E
F
G
H
E'
F'
G'
H'
E"
F"
G"
H"
Foxo3KO/KO
I
J
K
L
I'
J'
K'
L'
I"
J"
K"
L"

**Table Legend S3.**

**Supplementary Table 3. Gene clusters detected in comparison of *Foxo3*$^{KO/KO}$ versus WT cochleae at 4 HPN contain several stress response, calcium regulatory, and cytoskeletal factors.** Composition of genes identified during clustering analysis, listed with their membership percentage and function adapted from the GeneCards human gene database.

| ***Cluster 1*** | | |
|---|---|---|
| ***Genes*** | **Membership** | **Function** |
| *Atf3* | 72% | Stress responsive transcription factor |
| *Bhlhe40* | 82% | Transcription factor that represses expression of Per, a circadian rhythm gene |
| *Clcf1* | 81% | Potent neurotrophic factor that binds to the CNTF receptor |
| *Fosl1* | 72% | Leucine zipper protein that dimerizes with Jun to form AP1 |
| *Sdc4* | 80% | Transmembrane receptor |
| ***Cluster 2*** | | |
| ***Genes*** | **Membership** | **Function** |
| *Bspry* | 73% | May regulate epithelial calcium transport by inhibiting TRPV5 activity |
| *Epcam* | 84% | Homotypic calcium-independent cell adhesion molecule |
| *Itga5* | 75% | Integrin alpha chain family |
| *Krt18* | 86% | Encoded by RT18 |
| *Krt8* | 74% | Dimerizes with KRT18 to form an intermediate filament |
| *Runx1* | 73% | Transcription factor that forms a complex with CBFB |
| ***Cluster 3*** | | |
| ***Genes*** | **Membership** | **Function** |
| *Bach1* | 72% | Transcriptional regulator that represses transcription of genes under the the NFE2L2 oxidative stress pathway |
| *Ccnl1* | 80% | Involved in pre-mRNA splicing and functions in association with cyclin-dependent kinases (CDKs) |
| *Fam107b* | 73% | Protein coding gene |
| *Frat2* | 85% | Positive regulator of WNT signaling pathway |
| *Hilpda* | 84% | Increases intracellular lipid accumulation and stimulates expression of cytokines including IL6, MIF and VEGFA |
| *Nr4a3* | 82% | Transcriptional activator that can bind the NGFI-B Response Element (NBRE) |
| *Nuak2* | 71% | Stress-activated kinase involved in tolerance to glucose starvation. Induces cell-cell detachment by increasing F-actin conversion to G-actin. |
| *Pim1* | 76% | Involved in cell survival and proliferation; can phosphorylate and inhibit proapoptotic proteins BAD, MAP3K5, and FOXO3 |
| *Ppp1r15a* | 74% | Transcript levels are increased following stressful growth arrest conditions and treatment with DNA-damaging agents |
| *Rnf19b* | 77% | E3 ubiquitin-protein ligase that plays a role in the cytotoxic effects of natural killer (NK) cells |
| *Vcam1* | 71% | Mediates leukocyte-endothelial cell adhesion and signal transduction |
| ***Cluster 4*** | | |
| ***Genes*** | **Membership** | **Function** |
| *1700025G* | 91% | Unknown |
| *2610017I0* | 83% | Unknown |
| *2610035F* | 84% | Unknown |
| *4930402H* | 77% | Unknown |
| *5031439G* | 72% | Unknown |
| *Arhgap20* | 76% | Activator of RHO-type GTPases, transducing a signal from RAP1 to RHO and impacting neurite outgrowth |
| *Bcat2* | 73% | Branched chain aminotransferase found in mitochondria |

| | | |
|---|---|---|
| *Ccdc141* | 72% | Radial migration and centrosomal function |
| *Clmn* | 80% | Actin binding |
| *Crot* | 80% | Helps transport medium- and long- chain acyl-CoA molecules out of the peroxisome to the cytosol and mitochondria |
| *D630003M* | 82% | Unknown |
| *Epn2* | 75% | Clathrin-mediated endocytosis and Notch Signaling Pathway |
| *Esrrb* | 71% | Unknown but encodes a protein with similarity to the estrogen receptor |
| *Fat2* | 85% | Cell adhesion molecule |
| *G0s2* | 86% | Promotes apoptosis by binding to BCL2, preventing formation of protective BCL2-BAX heterodimers |
| *Gprc5b* | 93% | GPCR that may modulate insulin secretion and increased protein expression |
| *Itfg3* | 89% | Unknown |
| *Met* | 72% | Prototypical receptor tyrosine kinase |
| *Olfml1* | 71% | Unknown |
| *Pdk1* | 85% | Protects cells against apoptosis in response to hypoxia and oxidative stress |
| *Pou3f3* | 71% | Transcription factor that acts synergistically with SOX11 and SOX4 |
| *Ppp1r1b* | 72% | Dopaminergic and glutamatergic receptor stimulation regulates its phosphorylation and function as a kinase or phosphatase inhibitor |
| *Slc4a11* | 73% | Involved in transport of potassium through the fibrocyte layer to the stria vascularis and is essential for the generation of the endocochlear potential but not for regulation of potassium concentrations in the endolymph |
| *Sorcs3* | 72% | Type-I receptor transmembrane protein associated with Alzheimer's disease |
| *Tcea3* | 81% | Necessary for efficient RNA polymerase II transcription elongation past template-encoded arresting sites |
| *Tgfbr2* | 81% | Contributes to the transcription of genes related to cell proliferation, cell cycle arrest, and wound healing |
| *Tmtc4* | 72% | TPR domains mediate protein-protein interactions in various cellular processes |
| *Wnt5a* | 72% | Can activate or inhibit canonical Wnt signaling |
| *Zic2* | 92% | Transcriptional repressor that may regulate tissue specific expression of dopamine receptor D1 |
| ***Cluster 5*** | | |
| **Genes** | **Membership** | **Function** |
| *Igf1r* | 85% | Cell growth and survival control; can activate JNK kinases |
| *Pgm2l1* | 74% | Intramolecular transferase activity, phosphotransferases and glucose-1,6-bisphosphate synthase activity |
| *Sesn1* | 73% | Mediates p53 inhibition of cell growth by activating AMP-activated protein kinase, which results in the inhibition of the mammalian target of rapamycin protein |
| *Syt9* | 70% | May be involved in Ca(2+)-dependent exocytosis of secretory vesicles through Ca(2+) and phospholipid binding to the C2 domain or may serve as Ca(2+) sensors in the process of vesicular trafficking and exocytosis |
| *Tprgl* | 71% | Regulates synaptic release probability by decreasing the calcium sensitivity of release |

| | | |
|---|---|---|
| *Wdr45* | 81% | Component of the autophagy machinery that controls the major intracellular degradation process by which cytoplasmic materials are packaged into autophagosomes and delivered to lysosomes for degradation |
| *Ypel3* | 80% | Involved in proliferation and apoptosis in myeloid precursor cells |
| ***Cluster 6*** | | |
| **Genes** | **Membership** | **Function** |
| *Csrnp1* | 72% | Has transcriptional activator activity and may play a role in apoptosis. |
| *Dusp5* | 73% | Inactivates ERK1; negatively regulates members of the mitogen-activated protein (MAP) kinase superfamily (MAPK/ERK, SAPK/JNK, p38) |
| *Egr1* | 82% | Regulates gene transcription for cell survival, proliferation and cell death; can activate expression of p53/TP53 and TGFB1 |
| *Fos* | 76% | Dimerizes with proteins of the JUN family to form the transcription factor complex AP-1. |
| *Gadd45b* | 70% | Involved in the regulation of growth and apoptosis; mediates activation of stress-responsive MTK1/MEKK4 MAPKKK |
| *Gdf15* | 88% | Increased protein levels are associated with disease states such as tissue hypoxia, inflammation, acute injury and oxidative stress |
| *Gem* | 80% | Regulatory protein in receptor-mediated signal transduction |
| *Hbegf* | 86% | Growth factor that mediates its effects via EGFR, ERBB2 and ERBB4 |
| *Junb* | 71% | Transcription factor involved in regulating gene activity following the primary growth factor response |
| *Lif* | 80% | Pleiotropic cytokine |
| *Maff* | 80% | Involved in the cellular stress response |
| *Nfkbia* | 74% | Inhibits the activity of dimeric NF-kappa-B/REL complexes by trapping REL dimers in the cytoplasm through masking of their nuclear localization signals |
| *Phlda1* | 73% | Regulation of apoptosis |
| *Plk3* | 73% | Rapidly activated upon stress stimulation, such as ionizing radiation, reactive oxygen species (ROS), hyperosmotic stress, UV irradiation and hypoxia. |
| *Socs3* | 79% | Negative regulation of cytokines that signal through the JAK/STAT pathway |
| ***Cluster 7*** | | |
| **Genes** | **Membership** | **Function** |
| *Lamc2* | 88% | Mediates the attachment, migration and organization of cells into tissues during embryonic development |
| *Serinc2* | 78% | Viral mRNA Translation and Metabolism |
| *Socs2* | 72% | Regulates receptor turn-over; downregulates growth factors and upregualtes Trk receptors |
| ***Cluster 8*** | | |
| **Genes** | **Membership** | **Function** |
| *Aass* | 72% | Bifunctional enzyme that catalyzes the first two steps in lysine degradation |
| *Ahcyl2* | 72% | May be involved in the conversion of S-adenosyl-L-homocysteine to L-homocysteine and adenosine |
| *Appl2* | 87% | Multifunctional adapter protein and master regulator |

| | | |
|---|---|---|
| *Crygn* | 73% | Member of the crystallin family of proteins that are localized to the refractive structure of vertebrate eye lenses |
| *Gm19990* | 76% | Unknown |
| *Gpr30* | 81% | G-protein coupled estrogen receptor that binds to 17-beta-estradiol (E2) with high affinity, leading to rapid and transient activation of numerous intracellular signaling pathways |
| *Ksr1* | 70% | Part of a multiprotein signaling complex which promotes phosphorylation of Raf family members and activation of downstream MAP kinases |
| *Lctl* | 74% | Member of the glycosidase enzymes |
| *Maf* | 77% | Acts as a transcriptional activator or repressor |
| *Map2k6* | 81% | Dual specificity protein kinase which acts as an essential component of the MAP kinase signal transduction pathway |
| *Mapk4* | 92% | Phosphorylates microtubule-associated protein 2 (MAP2) and MAPKAPK5 |
| *Megf9* | 91% | Unknown |
| *Paqr8* | 87% | Unknown |
| *Rab17* | 73% | Epithelial cell-specific GTPase |
| *Rapgef3* | 72% | Guanine nucleotide exchange factor (GEF) for RAP1A and RAP2A small GTPases that is activated by binding cAMP |
| *Slc18b1* | 79% | Unknown |
| *Slc8a3* | 85% | Mediates the electrogenic exchange of Ca(2+) against Na(+) ions across the cell membrane, and thereby contributes to the regulation of cytoplasmic Ca(2+) levels and Ca(2+)-dependent cellular processes |
| *Tprkb* | 85% | Component of the EKC/KEOPS complex |
| *Tyr* | 71% | Catalyzes the initial and rate limiting step in the cascade of reactions leading to melanin production from tyrosine |
| *Wdr91* | 87% | Functions as a negative regulator of the PI3 kinase/PI3K activity associated with endosomal membranes via BECN1, a core subunit of the PI3K complex |

**Figure Legend S5.**

**Supplementary Figure 5. Eight clusters of genes with similar expression change patterns observed between *Foxo3*$^{KO/KO}$ versus WT cochleae.** Gene expression clustering was performed with Mfuzz for the *Foxo3*$^{KO/KO}$ versus WT cochleae at 4 HPN. DEGs had a fold change 2 threshold and false discovery rate (FDR) 5% correction. The resulting clusters contained genes with at least a 70% membership value. The expression rate (y-axis) is plotted against the time course of each genotype (x-axis). Clusters 4, 5, and 8 showed decreases in gene transcript detection for the *Foxo3*$^{KO/KO}$ at 4 HPN.

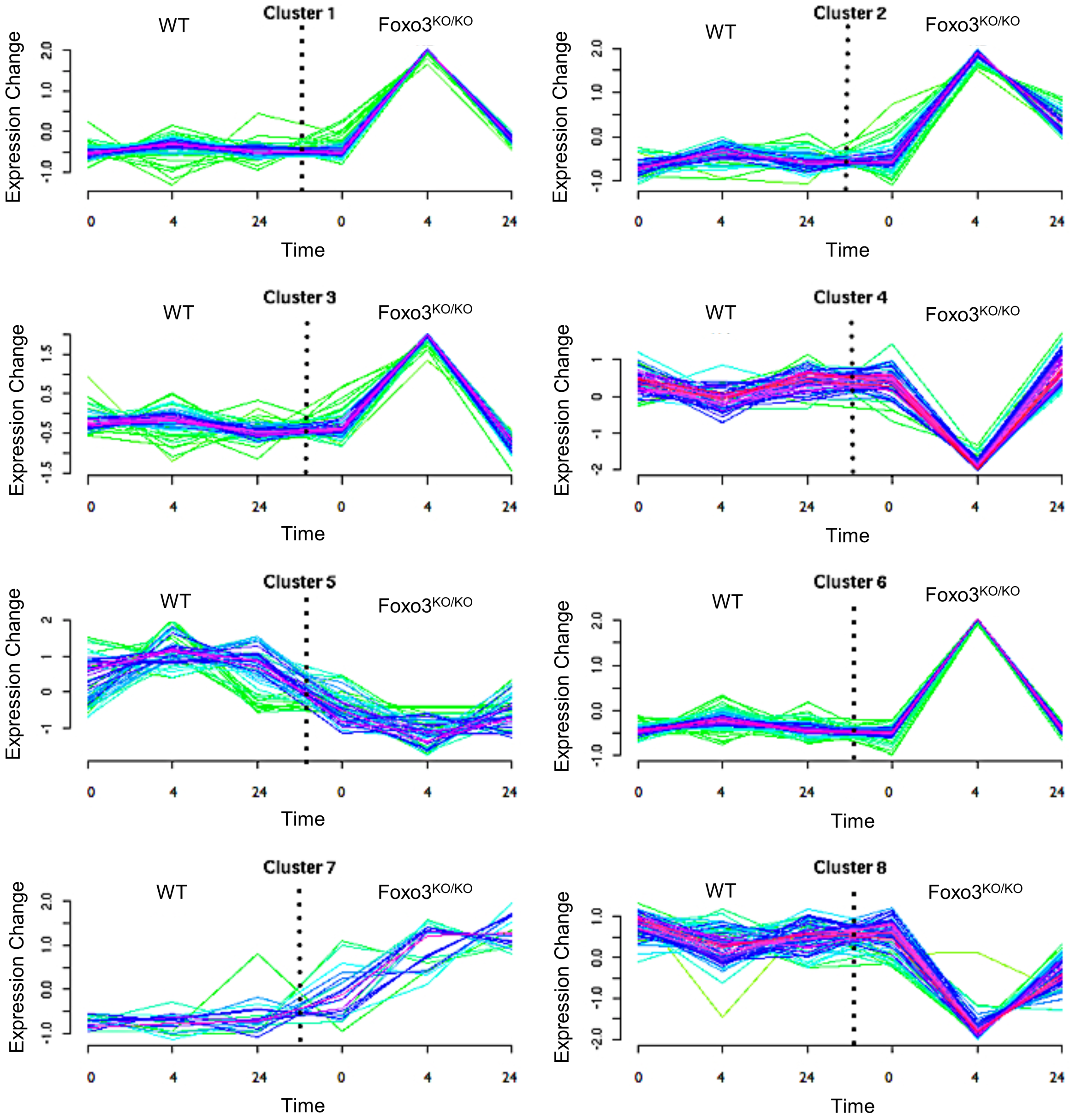

Cluster 1
WT
Foxo3$^{KO/KO}$
Expression Change
-1.0 0.0 1.0 2.0
0 4 24 0 4 24
Time
Cluster 2
WT
Foxo3$^{KO/KO}$
Expression Change
-1.0 0.0 1.0 2.0
0 4 24 0 4 24
Time
Cluster 3
WT
Foxo3$^{KO/KO}$
Expression Change
-1.5 -0.5 0.5 1.5
0 4 24 0 4 24
Time
Cluster 4
WT
Foxo3$^{KO/KO}$
Expression Change
-2 -1 0 1
0 4 24 0 4 24
Time
Cluster 5
WT
Foxo3$^{KO/KO}$
Expression Change
-1 0 1 2
0 4 24 0 4 24
Time
Cluster 6
WT
Foxo3$^{KO/KO}$
Expression Change
-1.0 0.0 1.0 2.0
0 4 24 0 4 24
Time
Cluster 7
WT
Foxo3$^{KO/KO}$
Expression Change
-1.0 0.0 1.0 2.0
0 4 24 0 4 24
Time
Cluster 8
WT
Foxo3$^{KO/KO}$
Expression Change
-2.0 -1.0 0.0 1.0
0 4 24 0 4 24
Time

**Figure Legend S6.**

**Supplementary Figure 6. No significant difference in Hsp70 expression within control OHCs and DCs between genotypes. A-C)** Single-matched optical sections from confocal images of 24 kHz OHCs under control conditions. WT **(A)**, *Foxo3*$^{+//KO}$ **(B)** and *Foxo3*$^{KO/KO}$ **(C)** cochleae were sectioned and immunostained for detection of Hsp70 (white) in OHCs and DCs. n = 3 per genotype, 60x magnification, scale bar = 10 μm. **D)** Quantification of the mean corrected total cell fluorescence (CTCF) of Hsp70 for OHCs and DCs. Each dot represents the CTCF means of each section and the bars are the means of those values: WT (black circles, orange mean bar), *Foxo3*$^{+//KO}$ (gray squares, pink mean bar), and *Foxo3*$^{KO/KO}$ (light gray triangles, green mean bar). No significant differences in Hsp70 expression were observed between genotypes. Two-way ANOVA with Tukey's test for multiple comparisons, alpha = 0.05.

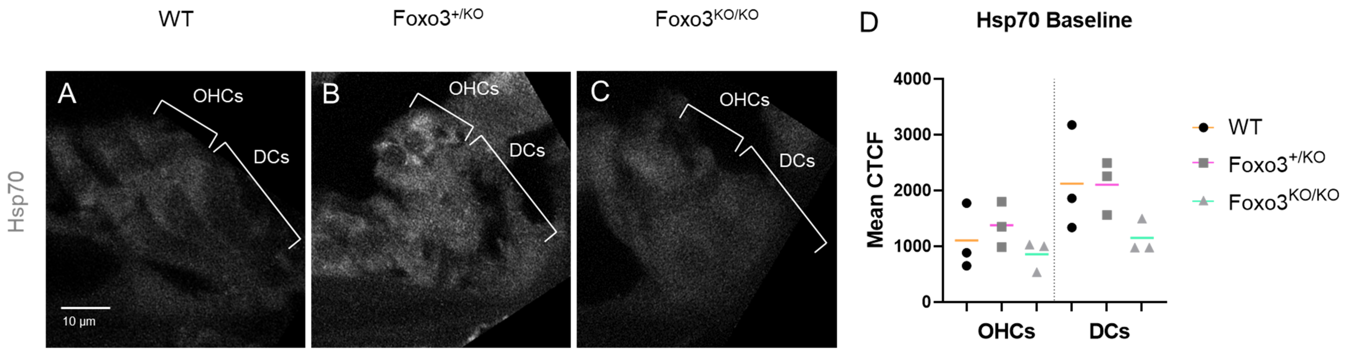
WT
Foxo3+/KO
Foxo3KO/KO
D
Hsp70 Baseline
A
OHCs
DCs
B
OHCs
DCs
C
OHCs
DCs
Hsp70
10 µm
4000
3000
2000
1000
0
Mean CTCF
OHCs
DCs
WT
Foxo3+/KO
Foxo3KO/KO

**Figure Legend S7.**

**Supplementary Figure 7. Expression of p53 is not strongly modulated following noise exposure.** WT and *Foxo3*$^{KO/KO}$ mice were exposed to noise and their cochleae were extracted for cryosectioning and immunohistochemistry at designated time points. **A-F')** Cochlear sections of the 24 kHz region immunostained for DAPI (cyan), OCM (magenta), and p53 (white) at baseline, 0.5 HPN, and 4 HPN. n = 3-5 per genotype/condition, 200x magnification, scale bar = 10 µm. Corresponding % fluorescent cellular expression graphs below each set of images: undetected = white, low = light gray, high = gray, and lost cells = black; DC: Deiters' Cells, OHCs: outer hair cells, OHC S&N: OHC synapses and neurites, PCs: pillar cells, IHC S&N: IHC synapses and neurites, and IHC: inner hair cells. With the exception of IHCs, weak p53 levels are detected in most cells of the organ of Corti with no strong changes seen in either genotype after noise.

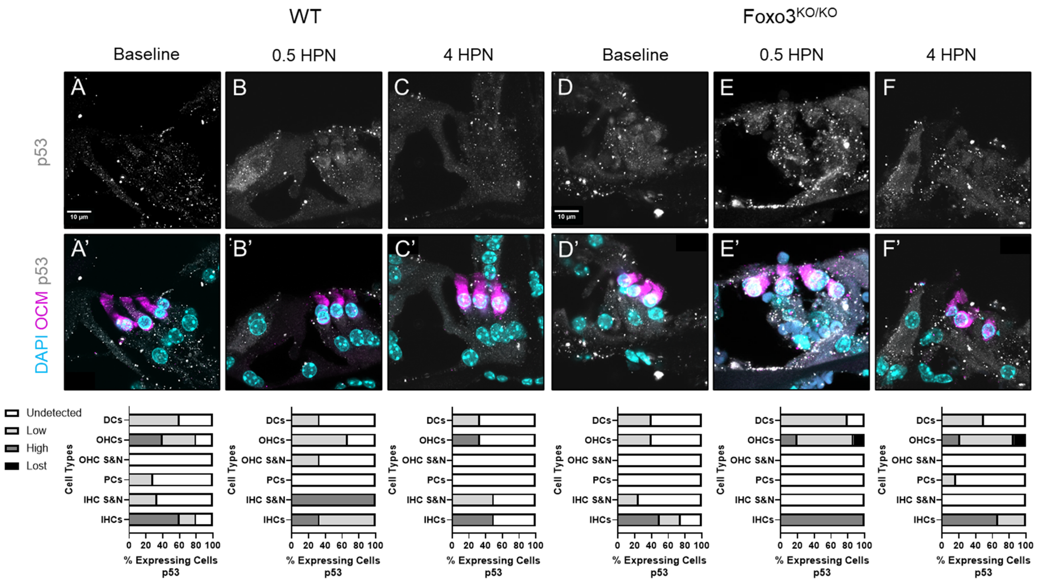

WT
Foxo3$^{KO/KO}$
Baseline
0.5 HPN
4 HPN
A
B
C
D
E
F
A'
B'
C'
D'
E'
F'
p53
DAPI OCM p53
10 μm
Undetected
Low
High
Lost
DCs
OHCs
OHC S&N
PCs
IHC S&N
IHCs
Cell Types
0 20 40 60 80 100
% Expressing Cells
p53

**Figure Legend S8.**

**Figure S8. GDPD3 is expressed in cochlear supporting cells and neurites. A, B)** Confocal images of GDPD3 (white) expressed in the pillar cells, Deiters' cells, and neurites of both WT and $\mathit{Foxo3}^{KO/KO}$ cochleae at baseline. DAPI (cyan), Parvalbumin (PARV, magenta), 24 kHz region, n = 3 per genotype, 60x magnification, scale bar = 20 μm.

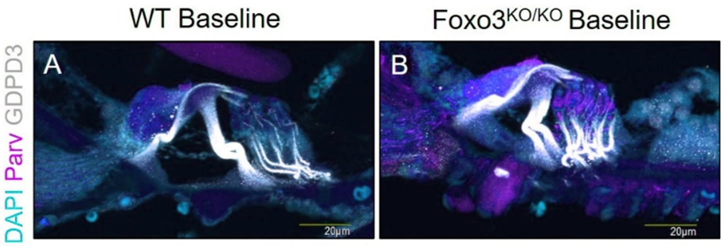
WT Baseline
Foxo3KO/KO Baseline
DAPI Parv GDPD3
A
B
20µm
20µm
